# Identification and deletion of the extracellular polysaccharide slime pathway across the genus Xanthobacter

**DOI:** 10.64898/2026.10.16.759691

**Authors:** Maximillian P.M. Soltysiak, Amogh P. Jalihal, Nathan R. Schaumburger, Michael Springer

## Abstract

Foods and commodities made by microbial bioproduction offer a sustainable alternative to those produced from our traditional petrochemical-based supply chains. Increasingly, *Xanthobacter* strains are used for bioproduction because of their unique metabolic properties (e.g., CO_2_ and N_2_ fixation) and wide feedstock range, including growth on agricultural products (e.g., fructose, sucrose) and electrically generated substrates (e.g., hydrogen gas, methanol, acetate, and formate). Many *Xanthobacter* strains, however, produce an extracellular polysaccharide (EPS) slime that makes plating, pelleting, and handling these cultures challenging and shuttles away resources from biomass and other biosynthetic pathways. While slime-free mutants of slime-producing *Xanthobacter* strains have been isolated, the causative mutations were unknown. Thus, slime-free mutants need to be painstakingly reisolated based on phenotypic properties in each academically and industrially relevant strain, unless we know the gene to target. Here, we identified the gene and underlying mutation that causes a slime-free phenotype in the *X. autotrophicus* mutant 7CSF. When we deleted this gene, *xsbA*, slime production was abolished in the wild-type strain. We showed that deleting *xsbA* homologs similarly abolished slime production in slime-producers across the genus *Xanthobacter* (*X. versatilis, X. autotrophicus, X. flavus,* and *X. tagetidis*). We also identified a Wzx/Wzy-dependent EPS pathway downstream of *xsbA* that is conserved across slime-producing *Xanthobacter* strains. Lastly, abolishing slime production improved colony formation and pelleting, with minimal impact on growth. We demonstrated that challenging phenotypic properties in *Xanthobacter* strains can be removed, improving their usability as a chassis for sustainable bioproduction.

**Importance:** *Xanthobacter* strains are promising as microbial platforms for bioproduction to more sustainably make our foods and commodities than current petrochemical-based methods. However, many of these strains produce a sugary slime that can make handling these strains difficult and that shuttles resources away from more desirable pathways. While slime-free mutants of *Xanthobacter* strains have been isolated before, the mutation that caused this change was unknown, and slime-free mutants would need to be reisolated each time. Here, we identify a gene, designated *xsbA*, that can be deleted in slime-producing *Xanthobacter* strains to remove their ability to make slime. Strains with the gene deleted become easier to handle and process, with minimal impact on growth. We also discovered additional genes involved in *Xanthobacter* slime production, which could be used to manipulate and engineer the properties of their slime for useful purposes.

## Introduction

Our world is on a dangerous climate trajectory due to our industrial activities and consumption. Global sustainable development goals have been set that, if met, would allow us to avoid most upcoming environmental and economic disasters due to climate change (1–3). But we are currently too reliant on petrochemicals to produce our food and commodities for us to meet these sustainability goals (4). Microbial bioproduction is an attractive path to circumvent these limitations because these platforms use land, water, and energy more efficiently (5–8). To capitalize on the potential of bioproduction, we need strains with good industrial properties.

*Xanthobacter* strains are a promising chassis for sustainable bioproduction in industry and academia because of their unique metabolic properties and resilience (9–16). Some strains have been grown at industrial scales and use a wide, flexible range of feedstocks, including agricultural products such as fructose and sucrose, and electrically generated substrates such as methanol, ethanol, acetate, and formate. Beyond organic substrates, *Xanthobacter* strains can grow chemolithoautotrophically on gaseous substrates (CO_2_, H_2_) and can fix nitrogen (N_2_). Exploiting these diverse metabolic capabilities, companies are already using *Xanthobacter* strains to produce sustainable fertilizers (15, 17) and protein (18, 19).

While *Xanthobacter* strains are promising as sustainable bioproduction platforms, a limitation is that many strains make an extracellular polysaccharide (EPS), often referred to as slime (9, 10, 20). EPS production has been industrially useful in other bacteria (e.g., xanthan gum from *Xanthomonas campestris* (21)), but in *Xanthobacter* strains it can impair growth and handling. Metabolically, the slime is a resource-rich product, made of chains of sugars (9, 22), which shuttles away carbon and energy from cellular biomass and other biosynthetic pathways. Physically, the slime complicates how we handle *Xanthobacter* strains. The slime can appear as a halo around colonies on a plate (described as a “fried egg” (9)) and or it can make colonies waxy, gooey, or swollen, often fusing them together (Fig. 1A). Liquid cultures of some strains are viscous (9), which complicates or prevents reliable pipetting and pelleting (Fig. 1B). Even strains that do not visually appear to produce slime can produce enough to interfere with centrifugal pelleting. While slime production is often growth dependent, the growth conditions that trigger EPS production are variable and depend on both the carbon substrate and the strain. As a result, across the genus *Xanthobacter*, there is no known growth condition that consistently eliminates slime production.

**Fig 1.**
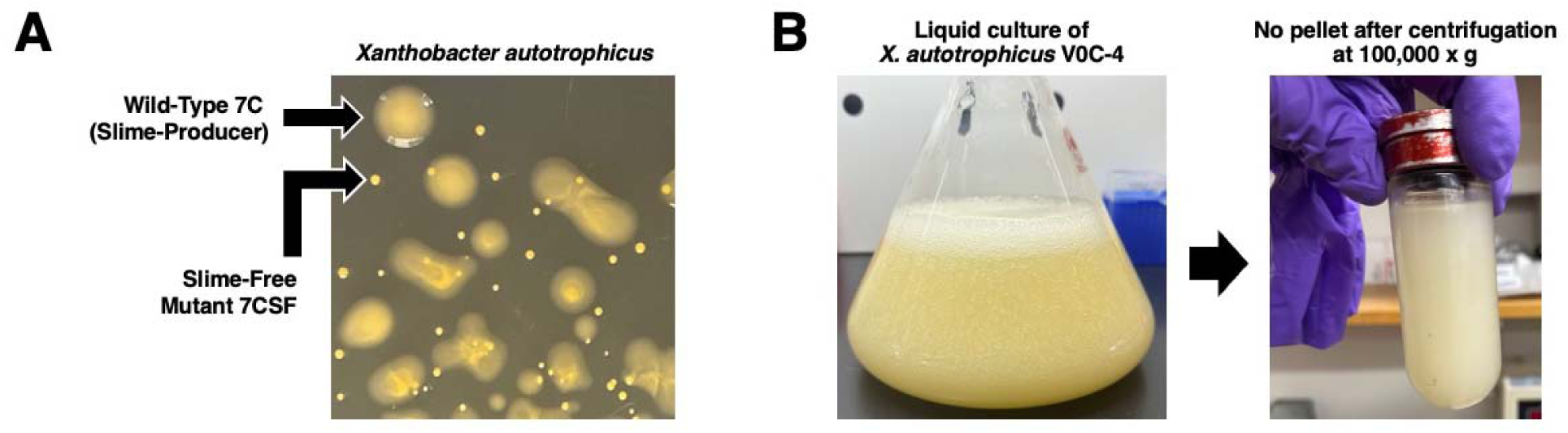
(A) Colonies of *X. autotrophicus* 7C (DSM 432) and its slime-free mutant 7CSF (DSM 2267) plated on rich media containing sucrose (a heavy slime-inducing condition). (B) Culture of slime-producer *X. autotrophicus* V0C-4 grown on minimal media containing 1.5% w/v β-hydroxybutyrate for 14 days. The strain produces copious amounts of slime, gelatinizing the culture and preventing pelleting even after centrifugation at 100,000 x *g* for 1 hour.

One solution to this problem would be to delete the slime production pathway. Slime-free mutants of slime-producing *Xanthobacter* strains have been isolated and described (23–25), but the causative mutations were unknown. Thus, slime-free mutants must be phenotypically reisolated in each strain. Identification of the genes necessary for slime production would greatly facilitate the reliable creation of slime-free phenotypes in academically and industrially relevant *Xanthobacter* strains.

In this study, we identified the gene (which we designated *xsbA*) and underlying mutation responsible for the loss of slime production in the previously described slime-free mutant *Xanthobacter autotrophicus* 7CSF. We demonstrated how homologs of this gene can be deleted in diverse slime-producers across the genus *Xanthobacter* to abolish slime production and improve handling. We also identified a conserved gene cluster related to EPS production in *Xanthobacter* strains, a target for future characterization, and a potential target for manipulating and engineering their slime properties.

## Results

### Identification of a slime-free mutation in Xanthobacter autotrophicus

To identify mutation(s) that can cause a slime-free phenotype in *Xanthobacter*, we ordered, sequenced, and compared the previously described slime-free mutant *X. autotrophicus* 7CSF (DSM 2267) to its wild-type parent 7C (DSM 432) (23). The highest-confidence mutation identified by the bcftools (26) variant-calling pipeline was in a gene annotated as an “undecaprenyl-phosphate glucose phosphotransferase” (Table S1). Homologs of this gene are involved in exopolysaccharide production in other model slime-producing bacteria, including *Ensifer/Rhizobium/Sinorhizobium meliloti* (exoY, succinoglycan production (27)) and *Xanthomonas campestris* (gumD, xanthan gum production (28–30)). In both species, this protein attaches the first sugar phosphate to a polyprenyl phosphate lipid carrier in the membrane during Wzx/Wzy-dependent exopolysaccharide biosynthesis (31, 32). When these initiating sugar phosphate transferases are disrupted or deleted in *Ensifer* and *Xanthomonas*, slime production is abolished. Thus, we hypothesized that *X. autotrophicus* 7C also relies on this initiating sugar phosphate transferase gene for slime production. After validating its role in slime production (below), we designated this gene as *xsbA* (“*<u>X</u>anthobacter* <u>s</u>lime <u>b</u>iosynthesis A”). The protein alignment of XsbA homologs from 7CSF, 7C, and other model slime-producers shows that a conserved proline residue is mutated in *X. autotrophicus* 7CSF (Fig. 2A), which may cause a loss of protein function and, ultimately, slime production.

**Fig 2.**
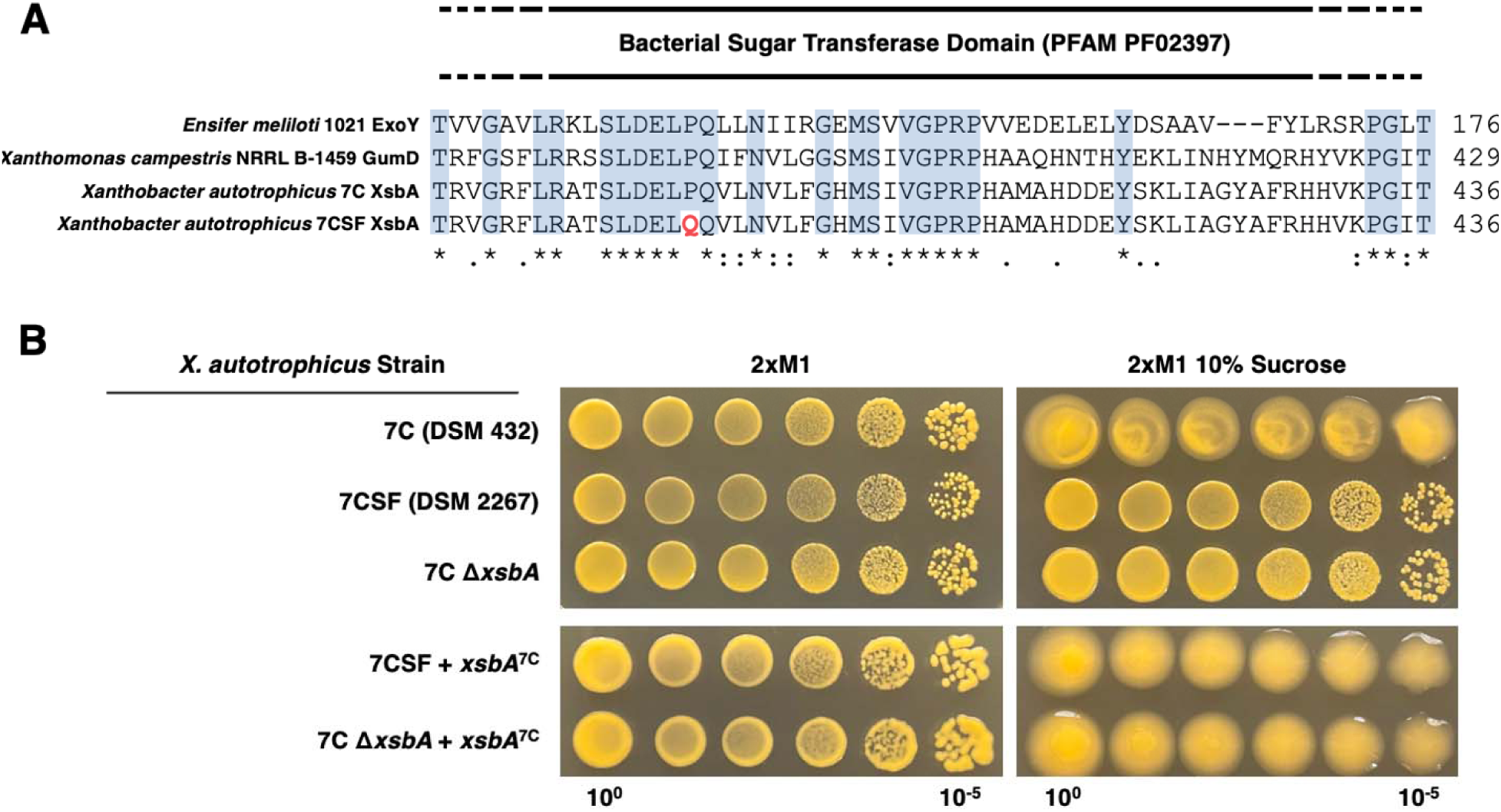
(A) Partial Clustal-Omega sequence alignment of the initiating polyprenyl phosphate sugar phosphate transferase homologs from *Xanthobacter autotrophicus* 7CSF (XsbA, WP_394025923.1) and its wild-type parent 7C (WP_138398082.1), and from model slime-producers *Ensifer meliloti* (ExoY, WP_010975907.1) and *Xanthomonas campestris (*gumD, WP_057672146.1). Conserved residues are highlighted in blue. Slime-free mutant *X. autotrophicus* 7CSF contains a mutation that changes a conserved proline residue to a glutamine (highlighted in red) within the bacterial sugar transferase domain (annotated by InterPro) of XsbA. Amino acid residue numbering of the protein sequences are shown on the right. (B) Deletion of *xsbA* abolishes slime production in *X. autotrophicus* 7C, an effect that is easier to observe under heavy slime-inducing conditions, such as when sucrose is in the media. Slime production can be recovered by introducing a wild-type copy of the 7C *xsbA* gene on a plasmid into slime-free mutant 7CSF and the slime-free deletion strain Δ*xsbA*.

To confirm that *xsbA* was needed for slime production in *X. autotrophicus*, we deleted *xsbA* in the wild-type parent 7C using seamless homologous recombination. Even under heavy slime-inducing conditions (i.e., sucrose), the Δ*xsbA* strain colonies were distinct and pinpoint, and produced minimal or no slime (Fig. 2B). We then expressed the wild-type *xsbA* (cloned from *X. autotrophicus* 7C) on a plasmid under its native promoter to see whether it was sufficient to restore slime production in 7CSF. Expression of wild-type 7C *xsbA* restored slime production in 7CSF to levels similar to those in the parent wild-type strain. Given the homology of *xsbA* to other initiating polyprenyl phosphate sugar phosphate transferases and the similarity in phenotypes when deleting this gene, it is very likely that *xsbA* performs the critical first step of slime production in *X. autotrophicus* 7C.

### Deleting xsbA to abolish slime production is a genus-wide solution for Xanthobacter strains

Many *Xanthobacter* strains produce slime, and the conditions in which they do so are dependent on the species, strain, and carbon source (Fig. S1). No universal slime-free growth condition is known. Thus, identifying and optimizing conditions that support fast, high-density growth while minimizing slime production takes substantial work in each strain. Knowing that slime production in *X. autotrophicus* 7C can be eliminated by deleting *xsbA*, we hypothesized that this strategy could be used in other slime-producing *Xanthobacter* strains. Using BLAST, we found that all the slime-producing strains from *Xanthobacter* Clades I, II, and IV (10) we analyzed contained a highly homologous copy of *xsbA*, with 74 to 99% nucleotide identity to strain 7C (Table S2, Fig. S2). Furthermore, each of these *xsbA* homologs were found in a nearly identical genomic context. In each case, *xsbA* was found directly or shortly downstream of the same three proteins as in strain 7C: an alpha-E superfamily protein, a circularly-permuted type 2 ATP-grasp protein, and a competence/damage-inducible protein/CinA (Fig. S3). While not directly involved in slime production, based on the functional annotations, these 3 genes are likely involved in glutathione biosynthesis and stress. Based on the conserved colocalization and high shared sequence identity, we expected that the *xsbA* homologs we identified were also needed for EPS production in the slime-producing *Xanthobacter* strains.

We directly tested whether the *xsbA* homologs are necessary for slime production in several well-studied slime-producing *Xanthobacter* strains by deleting their copy of *xsbA* using seamless homologous recombination. Strains tested included *X. versatilis* Py2 (Clade I), *X. autotrophicus* V0C-4 (Clade I), *X. flavus* GJ10 (Clade II), and *X. tagetidis* TagT2C (Clade IV). We plated a serial dilution of the wild-type and Δ*xsbA* versions of each strain on media containing sucrose (heavy slime-inducing condition) and observed colony morphology after 7 days. In each case, deleting *xsbA* caused a clear visual elimination of slime production (Fig. 3).

**Fig 3.**
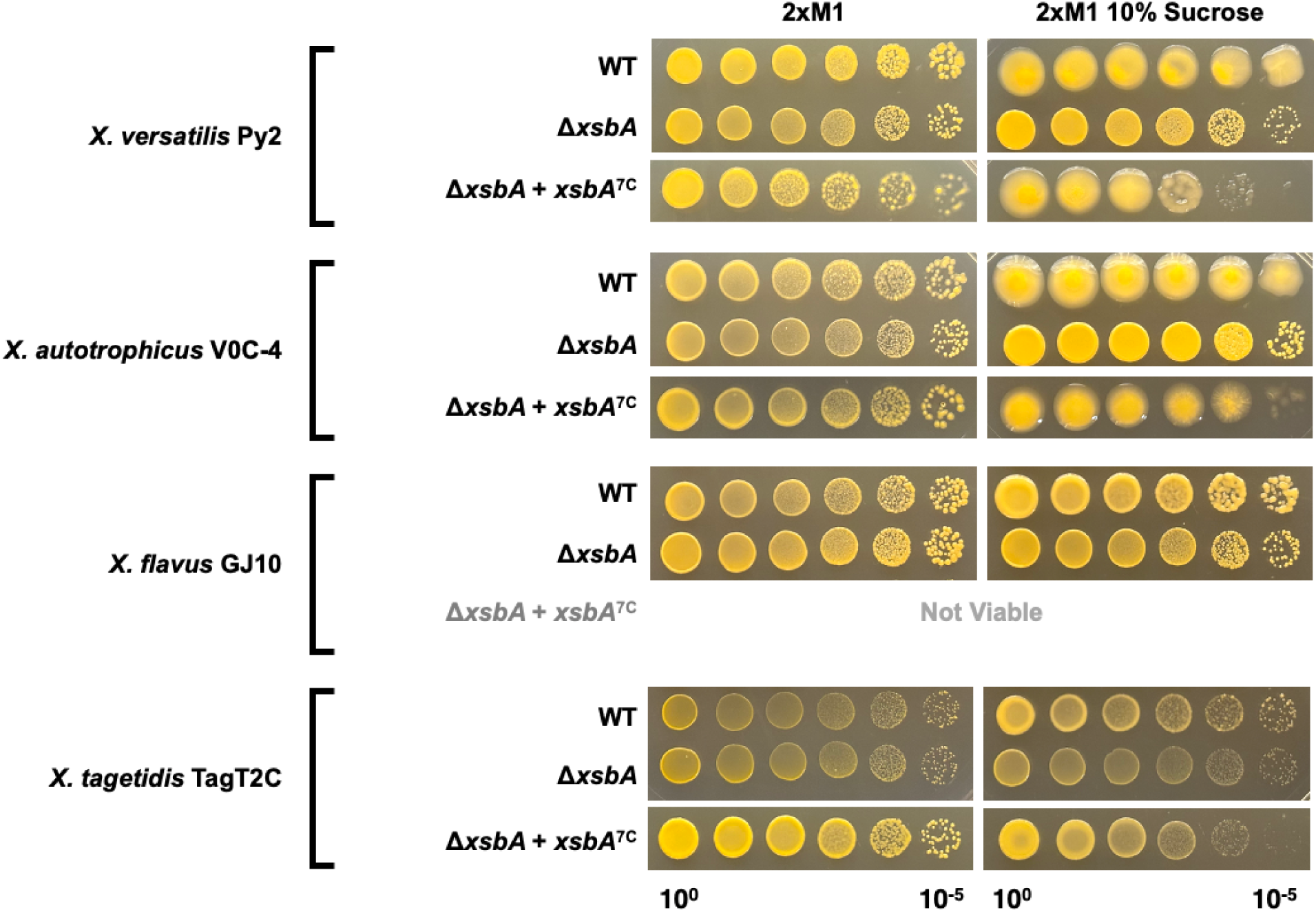
Deleting *xsbA* in diverse slime-producing *Xanthobacter* strains abolishes slime production. Slime production can be recovered in these Δ*xsbA* deletion strains by expressing wild-type *xsbA* from *X. autotrophicus* 7C, except in *X. flavus* GJ10.

To determine if the *xsbA* homologs perform the same interchangeable function, we tested whether the *X. autotrophicus* 7C *xsbA* gene could complement the *xsbA* deletion in the diverse strains and restore slime production. We expressed the 7C *xsbA* gene on a plasmid under its native promoter in each Δ*xsbA* strain and observed colony morphology after 7 days (Fig. 3). In the *X. versatilis* Py2*, X. autotrophicus* V0C-4, and *X. tagetidis* TagT2C strains, 7C *xsbA* could be transformed and successfully restored slime production. Even after multiple attempts, we were unable to transform the 7C *xsbA* plasmid into the waxy *X. flavus* GJ10, and hence we could not assess this hypothesis in this background. Regardless, we found that *xsbA* is a reliable target for deletion to abolish slime production in strains across the genus *Xanthobacter*.

### Identification of a Wzx/Wzy-dependent extracellular polysaccharide biosynthetic gene cluster in Xanthobacter autotrophicus

Slime-producing bacteria often possess many genes that alter the composition and structure of their EPS, usually found in a cluster or operon (27–29). We wanted to identify these other genes involved in *Xanthobacter* slime production. Many of these genes are likely annotated as “hypothetical proteins” or with vaguely related functional annotations, meaning a targeted gene deletion panel would miss many of them. A classic deletion screen looking for genes involved in slime production is also unlikely to be successful because in other model slime-producers, such as *Ensifer* and *Xanthomonas*, some intermediate genes in the Wzx/Wzy-dependent slime pathway could not be deleted in a wild-type background, likely due to the accumulation of toxic intermediates (29, 33). However, if the gene that produces the substrate for the initiating polyprenyl phosphate sugar phosphate transferase (i.e., the homolog of *xsbA*) is deleted in these species, downstream slime intermediates cannot accumulate as the initial lipid carrier for the EPS is not made. Thus, in these deletion backgrounds, the genes responsible for the enzymatic steps forming those toxic intermediates can now be deleted (27, 29). Following this logic, we hypothesized that some intermediate genes in the slime pathway in *Xanthobacter* could be deleted in a Δ*xsbA* background, but not in the wild-type background. To identify these genes, we created genome-wide gene deletion libraries using Tn5 transposon mutagenesis in both wild-type and Δ*xsbA* backgrounds of *X. autotrophicus* V0C-4 (Tables S3-7, Figs. S4-6, Fig. 4A). We selected V0C-4 as a model over 7C, as V0C-4 is highly transformable, making it easy to generate large numbers of colonies, each with a unique transposon insertion. We pooled and sequenced the two transposon libraries and looked for genes with higher insertion rates in the slime-free Δ*xsbA* background than in wild-type, as these would be candidates for intermediate genes in the slime pathway.

**Fig 4.**
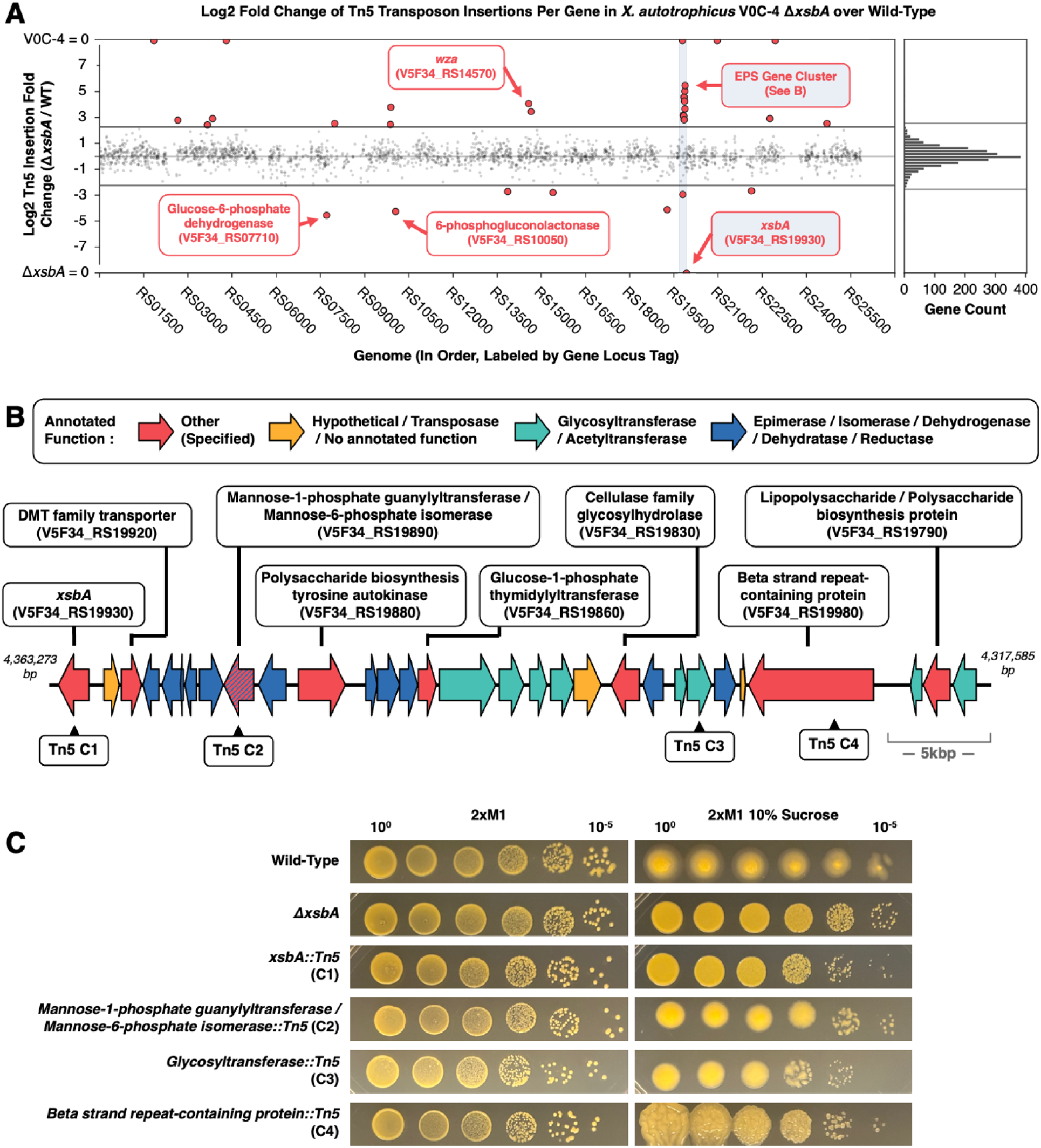
(A) Log2 fold change of Tn5 transposon insertions per gene in the *X. autotrophicus* V0C-4 Δ*xsbA* strain library over wild-type. Genes that had significantly different insertion counts between the two strain libraries are colored in red (p<0.05, Bonferroni corrected). A cluster of several genes with more insertions in the Δ*xsbA* background reveals likely intermediates in the EPS slime biosynthetic pathway (colored in blue), as well as a *wza* homolog elsewhere in the genome. Genes related to the pentose phosphate pathway are found with more insertions in the wild-type library than Δ*xsbA* library (6-phosphogluconolactonase and glucose-6-phosphate dehydrogenase). Genes with fewer than 10 Tn5 insertions in the wild-type library and Δ*xsbA* library have been filtered out (see Methods). The line at the top and bottom of the graph indicates genes with insertions in one background but 0 insertions in the other. (B) An approximately 44 kbp cluster of 30 genes with annotated functions related to Wzx/Wzy-dependent extracellular polysaccharide synthesis. (C) Colonies with abnormal slime production were picked from the wild-type transposon library and had insertions that mapped to the EPS slime gene cluster.

A cluster of genes downstream of *xsbA* contained 8 of the 18 significantly more inserted genes in the Δ*xsbA* background over wild-type (Fig. 4A, Table S6). This led to the discovery of an approximately 44 kbp gene cluster of 30 genes, most with annotated protein functions related to EPS biosynthesis (Fig. 4B, Table S8). For example, the cluster included sugar reductases, epimerases, isomerases, dehydrogenases, dehydratases, glycosyltransferases, and acetyltransferases. These functional annotations were supported by an additional BLAST search against the carbohydrate-active enzyme database (CAZymes (34) via ezCAZy (35)), which yielded hits for the glycosyltransferases (families 2 and 4) and a glycosyl hydrolase (Table S8). We carried out an additional BLAST of the V0C-4 slime gene cluster proteins against proteins from the Wzx/Wzy-dependent EPS pathways of *Ensifer meliloti* (succinoglycan; *exo* operon (27)), *Xanthomonas campestris* (xanthan; *gum* operon (28, 29)), and *Escherichia coli* (colanic acid; Wza, Wzc, Wzx, Wzy, and Wzz proteins (31, 36)). The *X. autotrophicus* V0C-4 slime cluster returned hits to some of the key proteins in those pathways, including Wzc, Wzz, ExoA, ExoB, ExoI, ExoN, ExoP, ExoQ, ExoY, GumD, GumI, and GumH (Table S8).

While a large number of the significantly more inserted genes in the Δ*xsbA* background were located within this gene cluster, others were found elsewhere in the genome. A gene annotated as a *wza* homolog was also identified during the transposon screen with 16-fold more insertions in the Δ*xsbA* background than in wild-type, consistent with the behavior expected for an intermediate gene involved in slime production (Fig. 4A, Table S6). While we wished to directly test whether all genes with significantly different insertion counts in the Δ*xsbA* background were directly involved in slime production, this would require extensive biochemical and genetic work, which was outside the scope of this manuscript. Additionally, while some genes in the slime pathway cannot be deleted unless *xsbA* is also deleted, it is likely that some intermediate genes still can be deleted in a wild-type background (and would not be identified by this screen). We therefore picked out and sequenced several transposon mutants from the wild-type V0C-4 library that had visually altered slime production to determine the site of their transposon insertion. Indeed, we found that some of these aberrant slime producers had transposon insertions in the *xsbA*-linked gene cluster (Fig. 4B and C). Thus, the genes within this cluster are likely directly involved in slime production. Together, the results from the transposon screen show that *X. autotrophicus* V0C-4 produces slime using a Wzx/Wzy-dependent pathway, with most genes colocalized within a cluster downstream of *xsbA*.

Some genes in the transposon screen behaved in the opposite way to what we expected of slime pathway intermediate enzymes, with significantly fewer insertions in the Δ*xsbA* background than wild-type (Fig. 4A, Table S7). One of these genes lies within the slime pathway gene cluster but does not have an annotated function. Elsewhere in the genome, two proteins with significantly fewer insertions in the Δ*xsbA* library were glucose-6-phosphate dehydrogenase and 6-phosphogluconolactonase. These proteins are involved in glucose-6-phosphate metabolism and catalyze the first two steps in shuttling glucose-6-phosphate into the pentose phosphate pathway (PPP) (37). Thus, the transposon screen also allowed us to identify other metabolic genes that are influenced by slime production.

### Core genes and syntenic structure of the slime pathway gene cluster are conserved in slime-producing Xanthobacter

We then asked whether the slime gene cluster identified in *X. autotrophicus* V0C-4 was conserved across other slime-producing *Xanthobacter* strains, both in gene content and gene order (synteny). We used clinker (38), an automated gene cluster analysis tool, to perform global all-vs-all alignments of the slime cluster proteins to identify homologous proteins (sequence identity over 30%) within the gene cluster across slime-producing strains (Fig. S3). Approximately a third of the V0C-4 gene cluster (10-12 genes) was conserved in slime-producers across the genus *Xanthobacter*, primarily the genes immediately downstream of *xsbA*. These conserved genes included a DMT family transporter; enzymes that modify dTDP-sugars and mannose; a reductase; and a likely Wzc protein (annotated as a polysaccharide biosynthesis tyrosine autokinase). Further downstream, slimy Clade I and IIA *Xanthobacter* (*X. versatilis*, *X. oligotrophicus*, *X. autotrophicus*, and *X. lutulentifluminis*) shared homologs and conserved syntenic structure with nearly the entire V0C-4 gene cluster.

Outside of Clade I, however, waxy Clade IIB *Xanthobacter* (*X. flavus, X. aminoxidans, X. wiegelii*) possessed different genes downstream of the 10 to 12 gene core, including other enzymes that modify GDP-mannose and GDP-fucose, which were absent from slimy *Xanthobacter* strains. Strains more distantly related to V0C-4 (e.g., Clade IV) lost most of the conserved genes and syntenic structure but still possessed a few colocalized homologs from the cluster. Meanwhile, *X. nonsaccharivorans* 14g had half of its pathway located elsewhere in the genome, not adjacent to *xsbA*, unlike in its close relatives. Additional differences appeared at the individual strain level, where some strains possessed unique hypothetical, polysaccharide-related, and glycosyltransferase proteins not found in V0C-4 or other strains. Meanwhile, other strains like *X. flavus* GJ10 and *Xanthobacter sp.* T101 had their gene clusters interrupted with transposable and other foreign elements. These differences all likely contribute to the wide range of slime phenotypes observed across the genus (Fig. S1). While gene content and syntenic structure of these EPS gene clusters changed in strains across the *Xanthobacter* genus, given the relevant annotations of the genes belonging to these clusters and colocalization with *xsbA,* they likely play a role in slime production in these strains.

### Abolishing slime production in Xanthobacter strains improves handling with minimal impact to growth

The goal of identifying a slime-free mutation or deletion was to improve the handling of slime-producing *Xanthobacter* strains. When we deleted *xsbA* in slime-producing *Xanthobacter* strains, colonies became more separable and pinpoint (Fig. 2B and 3). The deletion made single colonies easier to pick than wild-type strains, especially at higher cell densities. Additionally, cells of the slime-free strains became easier to pellet. We grew wild-type and Δ*xsbA* cultures of our representative *Xanthobacter* strains in a minimal slime-inducing medium (2xM1 rich media), a moderate slime-inducing medium (R2A) and in a heavy slime-inducing medium (minimal medium with 1.5% w/v β-hydroxybutyrate) and measured how they pelleted at 5,000 x *g* for 20 minutes, a speed and time common for routine laboratory handling (Fig. 5, Fig. S7). Wild-type strains of *X. versatilis* Py2 and *X. autotrophicus* V0C-4 pelleted well under minimal slime conditions, while *X. autotrophicus* 7C, *X. flavus* GJ10, and *X. tagetidis* TagT2C did not and had flaky or fluffy pellets. Under heavy slime-inducing conditions, however, no wild-type strain pelleted. In contrast, all Δ*xsbA* strains pelleted well, regardless of growth media. This is a considerable improvement given that some strains, like wild-type *X. autotrophicus* V0C-4, cannot be pelleted under heavy slime-inducing conditions even after centrifugation at 100,000 x *g* (Fig. 1B), but the slime-free Δ*xsbA* strain can readily pellet at 5,000 x *g* (Fig. 5). Thus, deleting *xsbA* in slime-producing *Xanthobacter* strains also improves the handling of their liquid cultures.

**Fig 5.**
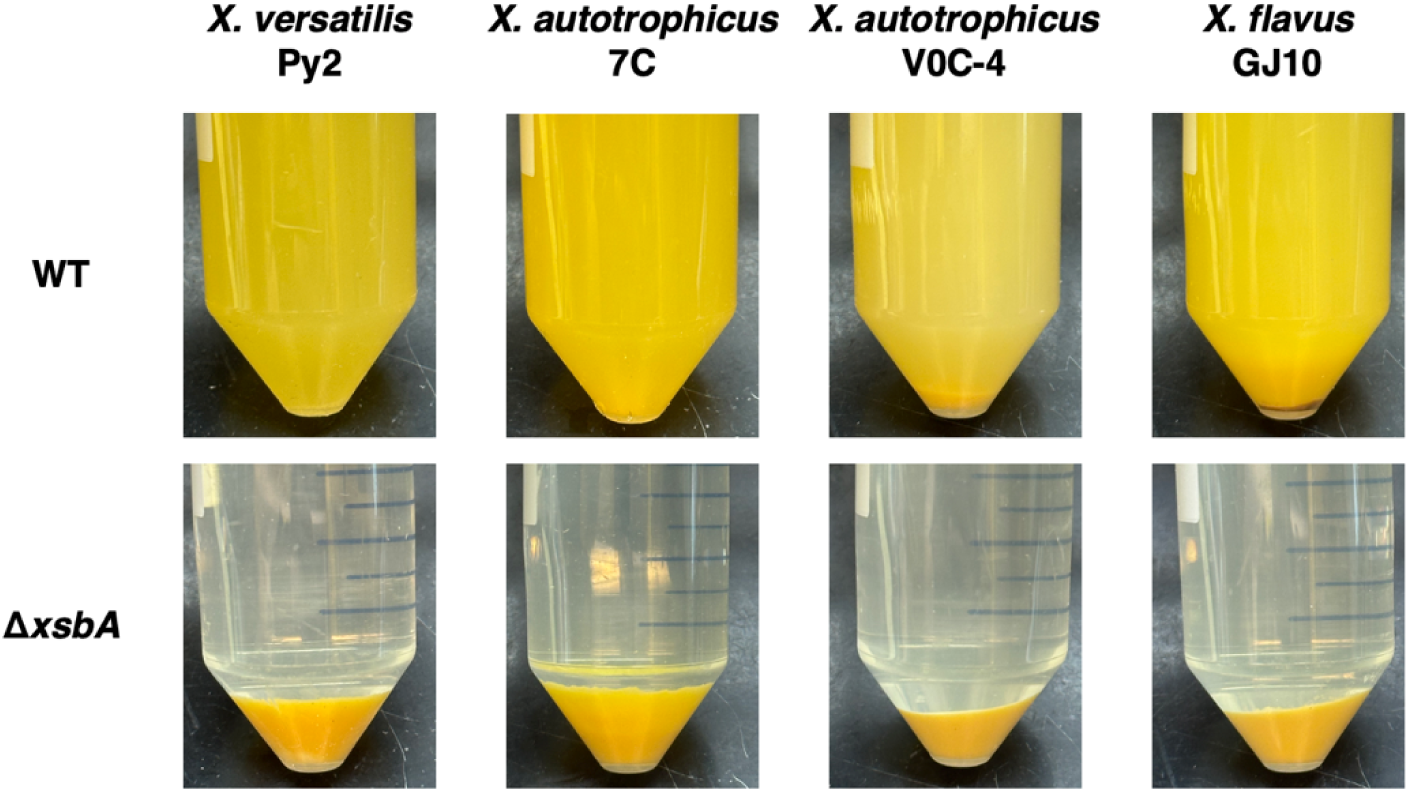
Slime-free Δ*xsbA Xanthobacter* strains pellet effectively, unlike their wild-type slime-producing equivalents. Cultures were grown under a heavy slime-inducing condition (1.5% w/v β-hydroxybutyrate) for 7 days, then spun down at 5,000 x g for 20 minutes.

Lastly, while handling is improved with the Δ*xsbA* deletion in slime-producing *Xanthobacter* strains, we wanted to confirm that these improvements did not come at the cost of growth rate and yield. We grew wild-type and Δ*xsbA X. autotrophicus* V0C-4 under two rich media conditions (2xM1 and R2A) and on three carbon sources (acetate, fructose, and sucrose). In each case, the slime-free strain grew as well as or slightly better than wild-type in terms of growth rate and density (Fig. 6). Additionally, we grew all wild-type and Δ*xsbA* strains of our representatives under fully autotrophic, nitrogen-fixing conditions (Fig. 6, Figs. S8 and S9). While measurements are noisier due to slight differences in gas delivery per culture, we did not observe a noticeable difference in growth rate or density outside of noise between the Δ*xsbA* and wild-type strains. Thus, abolishing slime production had minimal impact on the growth of slime-producing *Xanthobacter* cultures.

**Fig 6.**
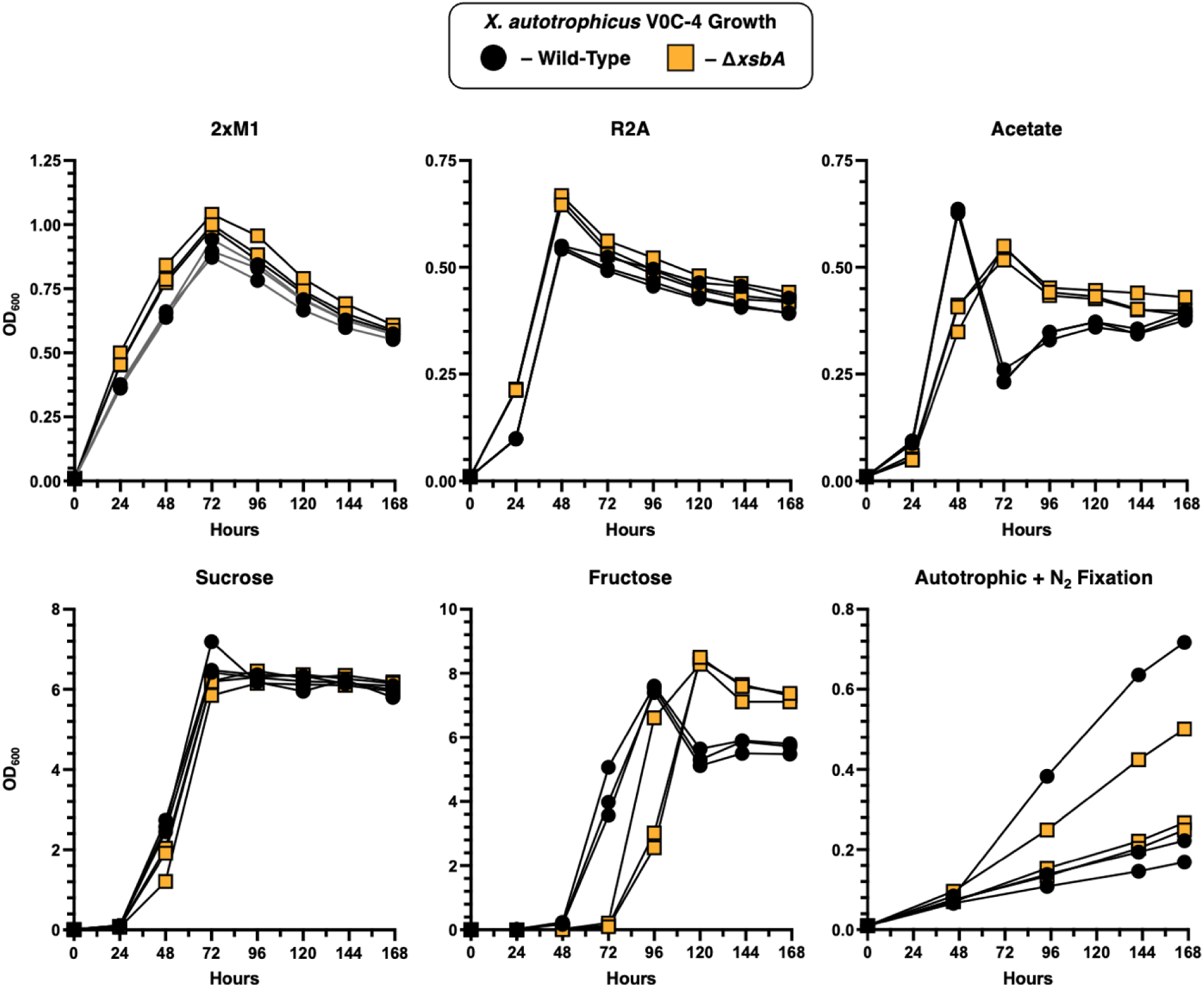
Optical density (OD_600_) measurements from liquid cultures of *X. autotrophicus* V0C-4 wild-type and Δ*xsbA* strains grown in either rich media (2xM1 and R2A), minimal media (with 0.5% w/v acetate, fructose, or sucrose), or in inorganic minimal media lacking a carbon or nitrogen source (bubbled with an autotrophic gas mixture: ∼2% O_2_, ∼3% H_2_, ∼10% CO_2_, rest N_2_).

## Discussion

We sought to identify a gene to target in slime-producing *Xanthobacter* strains that would allow us to abolish slime production and thus improve how we grow and handle these strains. Using an available slime-free mutant/wild-type pair from a repository (*X. autotrophicus* 7CSF and 7C) (23) we identified and found that deleting a gene annotated as an undecaprenyl-phosphate glucose phosphotransferase, designated *xsbA*, abolished slime production in *Xanthobacter* strains. Given that homologs of XsbA perform the first step in Wzx/Wzy-dependent EPS biosynthesis in other organisms (27, 29, 31), we reasoned that this gene is a good general target, as blocking the first committed step of the pathway, rather than one downstream, would prevent the buildup of potentially toxic intermediates (29, 33).

Variation in *xsbA* genes across different *Xanthobacter* strains could explain some of the differences seen in slime production between strains. For example, *xsbA* from *X. autotrophicus* 7C could complement *xsbA* deletions in slimy Clade I and Clade IV *Xanthobacter* but could not be transformed into the waxy *X. flavus* GJ10 (Clade II). This lack of complementation may indicate differences in the slime biosynthetic pathway between waxy strains (such as *X. flavus,* Clade II) and slimy/halo strains (such as *X. autotrophicus*, Clade I). The native *X. flavus* GJ10 *xsbA* might have evolved distinctly in this background and utilize a different sugar, or the protein might be expressed at a different level.

Differences in slime production and behavior across the genus are also likely due to the presence and absence of other genes outside of *xsbA*. In other organisms, there are many genes involved in EPS biosynthesis, usually localized in a cluster or operon. These genes influence the structure, composition, and properties of the EPS (27–29, 31). As expected, using a transposon screen, we found a slime biosynthetic gene cluster in the slime-producing *X. autotrophicus* V0C-4, with many functional annotations related to EPS production. Within the cluster there were numerous accessory proteins, including glycosyltransferases, acetyltransferases, reductases, dehydrogenases, and isomerases. But when we checked for core Wzx/Wzy-dependent pathway proteins from other model slime-producers, other than a likely Wzc homolog, we could not find obvious hits for the others. This is likely due to these core proteins notoriously lacking sequence conservation (31). Several proteins within the cluster had hypothetical functions or generic “lipopolysaccharide/polysaccharide biosynthesis protein” annotations and might be the missing core proteins and good targets for further characterization. Additionally, while these core proteins usually are found within slime operons/clusters of other organisms (27, 29, 31), they may still be elsewhere in the genome. For example, an annotated Wza homolog distant in the genome was a significant hit from our transposon screen, and likely was involved in the slime production pathway we were targeting. Regardless, given that nearly half of our hits from our transposon screen were within this cluster, their related functional annotations to EPS production, and behavior when knocked out, this gene cluster likely is the main driver of slime production in *X. autotrophicus* V0C-4. Furthermore, the core of the slime gene cluster was largely conserved in slime-producing strains across the genus *Xanthobacter.* The additional variation in gene content and syntenic structure we see likely explains differences in slime production, properties, and behavior between slime-producing *Xanthobacter* strains. Altering the composition of these gene clusters, including adding or removing unique proteins from other strains, could potentially be used to manipulate and engineer the structure and properties of *Xanthobacter* slime (39).

While genes with more insertions in the Δ*xsbA* transposon library than in wild-type were likely intermediate genes in the slime pathway, genes with the opposite behavior could also shed light on slime metabolism. For example, we saw more insertions in the wild-type background than Δ*xsbA* for two genes that shuttle glucose-6-phosphate into the pentose phosphate pathway (PPP) (37): glucose-6-phosphate dehydrogenase and 6-phosphogluconolactonase. The *X. autotrophicus* V0C-4 strain, however, cannot grow on exogenously supplied glucose. Yet it has been documented that *X. autotrophicus* slime contains glucose among its monomers (9, 22). Together, these facts suggest that EPS production in *X. autotrophicus* could be used as a metabolic shunt for eliminating glucose. When slime production is eliminated (i.e., *xsbA* is deleted), V0C-4 must rely on alternative pathways to deal with glucose accumulation and toxicity, namely the shunt into the PPP. If deleting the slime pathway in a *Xanthobacter* strain leads to poorer growth than in wild-type, the screen above suggests that engineering alternative metabolic shunts for slime monomers could improve growth or tolerance to the genetic modification.

Our motivation for this study was to identify a way to abolish slime production in *Xanthobacter* strains, as the slime complicated how we handle and grow these strains. When we deleted *xsbA* in slime-producing *Xanthobacter* strains, we saw improved colony formation and better pelleting. In these experiments, the discrepancies in pelleting between wild-type *Xanthobacter* strains under minimal slime-producing conditions raise some questions regarding the nature of their EPS. Some slimy strains, like *X. autotrophicus* V0C-4 and *X. versatilis* Py2, produce visible slime even under “minimal” slime conditions (i.e., a visible halo around colonies can be seen when grown on rich 2xM1 media, Fig. S1). These two strains, however, pellet well when grown on 2xM1. Yet, waxy strains (like *X. flavus* GJ10) and some other slimy strains (*X. autotrophicus* 7C, *X. tagetidis* TagT2C) do not appear to produce much or any slime under minimal slime-inducing conditions (including 2xM1). These strains, however, still do not pellet well when grown on 2xM1, with the loose pellets that form often appearing fluffy or flaky. In other bacteria, poor pelleting can be associated with extracellular polysaccharides covalently linked to the membrane, called capsular polysaccharides (40, 41). For the constitutive slime producers *X. autotrophicus* V0C-4 and *X. versatilis* Py2, since they still pellet well even when some slime is produced, this may indicate that their slime is not covalently linked to the membrane and is instead an exopolysaccharide. In practice, the distinction between exopolysaccharides and capsular polysaccharides is often unclear as it is difficult to measure (42), and because polysaccharides may or may not be covalently linked under different conditions in the same strain (43, 44). Overall, the reasons why different slime-producing *Xanthobacter* strains fail to pellet may differ between strains, including the supernatant having increased viscosity and density versus slime being covalently-linked to the outer membrane (9, 40). Regardless, deleting *xsbA* prevents slime production and allows for easy pelleting and colony picking of *Xanthobacter* strains.

In summary, we found that deleting *xsbA* in *X. autotrophicus* abolished slime production, and this effect was reproducible in other slime-producing *Xanthobacter* strains across the genus (*X. versatilis*, *X. flavus*, and *X. tagetidis*). Deleting *xsbA* improved the handling of these slime-producing strains, including colony formation and pelleting, with minimal or no impact to growth. We also identified a conserved gene cluster that drives slime production across the genus, an attractive target for further characterization and for manipulating the properties and composition of their slime. These findings show that we can reliably manipulate the phenotypic properties of *Xanthobacter* strains and improve their use as a bioproduction chassis.

## Materials and Methods

### Growth and Handling

*Xanthobacter* strains that were used in this research were isolated as described or sourced from the culture collections listed in Table S9. Cultures were grown as indicated in several media. 2xM1 medium contains: 10 g peptone (Gibco Bacto Peptone, Cat No: 211677) and 6 g meat extract powder (Gibco Difco Beef Extract Powder, Cat No: 212303) per L of Milli-Q water and was sterilized by autoclave. R2A medium (DSMZ Medium 830) contains: 0.5 g yeast extract (Gibco Bacto Yeast Extract, Cat No: 212750), 0.5 g proteose peptone (Gibco Bacto Proteose Peptone No. 3, Cat No: 211693), 0.5 g casamino acids (Gibco Bacto Casamino Acids, Cat No: 223050), 0.5 g glucose (Sigma-Aldrich, Cat No: G7021-100G), 0.5 g soluble starch (Sigma-Aldrich, Cat No: S9765-100G), 0.3 g sodium pyruvate (Alfa Aesar, Cat No: A11148), 0.3 g K_2_HPO_4_ and 0.05 g MgSO_4_ • 7 H_2_O per L of Milli-Q water and was sterilized by autoclave. The minimal medium used was modified Wiegel’s minimal medium (WMM)(45) which contains: 5.7 g K_2_HPO_4_, 1.95 g KH_2_PO_4_, 3 g (NH_4_)SO_4_, 0.2 g MgSO_4_ • 7 H2O, 0.04 g CaCl_2_ • 2 H_2_O, 0.03 g FeSO_4_ • 7 H_2_O and was sterilized by either filter or autoclave. After sterilization, 10 ml of a filter sterilized trace metal solution (modified from ATCC MD-TMS: 3 g MgSO_4_ • 7 H_2_O, 0.5 g MnSO_4_ • H_2_O, 1 g NaCl, 0.1 g FeSO_4_ • 7 H_2_O, 0.18 g CoCl_2_ • 6 H_2_O, 0.1 g CaCl_2_ • 2 H_2_O, 0.18 g ZnSO_4_ •7 H_2_O, 0.01 g CuSO_4_ • 5 H_2_O, 0.01 g H_3_BO_3_, 0.01 g Na_2_MoO_4_ • 2 H_2_O, 0.02 g NiCl_2_ • 6 H_2_O, 0.01 g Na_2_SeO_4_ dissolved in 1 L of pH 1 (using H_2_SO_4_) Milli-Q water) and 10 ml of a vitamin solution (ATCC Cat No. MD-VS) were added per L of modified WMM. The inorganic minimal medium (IMM) used contained: 1 g K_2_HPO4, 0.5 g KH_2_PO_4_, 2 g NaHCO_3_, 0.1 g MgSO_4_ · 7 H_2_O, 20 mL of a pre-dissolved 2 g/L CaSO_4_·2H_2_O solution, 0.01 g FeSO_4_ · 7 H_2_O per L of Milli-Q water. To this, 10 mL of the vitamin solution described above and 10 mL of the trace metal solution described above was added. For the autotrophic and nitrogen fixing plate experiments (Fig. S9), where indicated, WMM (lacking (NH_4_)SO_4_) and IMM plates did or did not include 2 g/L NaHCO_3_. If nitrogen was indicated, 3 g/L of (NH_4_)SO_4_ was added, and if carbon was indicated, 0.5% w/v di-sodium succinate was added. The LB media used for growing cloning and conjugative *E. coli* described below contained 10 g NaCl, 10 g peptone, and 5 g yeast extract. Agar plates of any of the media above contained 15 g agar (Gibco Difco Agar, Cat No: 214010) per L media and were sterilized by autoclave.

For Fig. S1, carbon substrates were added at 0.5% v/v (for alcohols) or 0.5% w/v (all others). For the pelleting experiment, 1.5% w/v β-hydroxybutyrate was used in the modified WMM. For heavy slime inducing conditions involving 2xM1 medium, 10% w/v sucrose (VWR BDH Chemicals Sucrose, Cat No: BDH9308-2.5KG) was added to the medium as well.

Xanthobacter cultures were grown at 30°C in either a stationary incubator (plates) or at 200 to 225 rpm in a shaking incubator (tubes, flasks). The OD_600_ values were measured using a spectrophotometer. Cultures were grown in 5 mL volumes in tubes prior to plating. For the pelleting experiments (Fig. 5 and Fig. S7), cultures were first grown in 20 mL 2xM1 to saturation prior to a 1:100 dilution into 50 mL volumes in 250 mL baffled flasks for growth for 7 more days in the media indicated. For the growth rate experiments (Fig. 6), saturated 2xM1 cultures were passaged 1:100 into 50 mL volumes in 250 mL baffled flasks in the new media indicated and grown for 3 days. These 3-day cultures were then passaged and restarted at an OD_600_ of 0.01 in the same fresh media indicated in Fig. 6 for the experiment shown. Pelleting was performed by centrifugation using 40 mL of culture at 5,000 x *g* for 20 minutes. The culture for Fig. 1B was grown in 2 L of 1.5% β-hydroxybutyrate modified WMM for 14 days. For Fig. 1B, culture was pelleted at 100,000 x *g* for 1 hour. For Fig. 1A, saturated 2xM1 cultures of *X. autotrophicus* 7C (DSM 432) and *X. autotrophicus* 7CSF (DSM 2267) were mixed 50:50 and a serial dilution was plated onto 2xM1 10% sucrose plates and grown for 7 days. A dilution where single colonies of each strain were visible was chosen for visualization.

Spot plating was done using 10-fold serial dilutions with 2xM1 medium prior to plating for all spot plating figures unless otherwise indicated. Spot plating with the plasmid expressing *X. autotrophicus* 7C *xsbA* was done on plates that included kanamycin (200 μg/mL for *X. tagetidis* TagT2C, 30 μg/mL for all others). Slime production was analyzed visually. We attempted to use the Congo Red Assay for exopolysaccharide production(46, 47) (Fig. S7), where 0.08% w/v of Congo Red Dye (Sigma-Aldrich, Cat No: C6277-25G) was added to the agar media, however, staining/coloration was variable across strains, and ultimately was not more informative than observing changes in sliminess by eye.

For autotrophic conditions, 30 mL cultures of wild-type and Δ*xsbA* strains were grown in a modified eVOLVER vial(10) at 30 °C with max spinning, with gases continuously bubbled in at the following concentrations: ∼2% O_2_, ∼3% H_2_, ∼10% CO_2_, and rest N_2_. Saturated 2xM1 cultures were passaged and first grown up in IMM with the autotrophic gases for 10 days prior to being passaged and restarted at OD_600_ of 0.01 for the experiment shown in Fig. 6 and Fig. S8. For the *X. tagetidis* strains, the autotrophic experiment was directly passaged from a 2xM1 culture instead. For the autotrophic and nitrogen fixing spot plates, cultures were first grown up in 2xM1, spun down as described above, and resuspended in IMM lacking nitrogen and carbon to an OD_600_ of 1.0, prior to a serial dilution for spot plating.

### Non-Transposon Bioinformatic Analyses

Short read sequencing reads for *X. autotrophicus* 7CSF were taken from our prior work where we published the genomes of most available *Xanthobacter* strains (10) (SRR40972798). Mutations in *X. autotrophicus* 7CSF compared to 7C were found by aligning the 7CSF reads to the *X. autotrophicus* 7C genome using bowtie2 (48) v2.5.1 (default settings), then samtools v1.17 sort, then bcftools (26) v1.17 mpileup (-d 1000, --min-BQ 20), bcftools call (--ploidy 1, -m, -v), and bcftools filter (-s LowQual -e ‘QUAL<20 || DP <10’) to generate the final table for Supplementary Table 1 (where mutations with a quality score above 100 are shown). An alignment of XsbA homologs was performed using Clustal-Omega (49) v1.2.4 via the EBI-EMBL webserver(50) (Fig. 2A, Fig. S2). Protein domains were annotated using InterPro(51) v109.0 through iprscan5 via the webserver (Fig. 1B) or InterPro v110 through iprscan6 (Fig. S2). BLAST analyses were performed using either blastn or blastp with default settings in Geneious v2026.0. For Supplementary Table 2, *X. autotrophicus* 7C *xsbA* was used as a query against custom BLAST (52) libraries made from all NT and AA sequences from the *Xanthobacter* representative/type genomes listed in that table. For Table S8, a custom BLAST database was made using the V0C-4 cluster proteins and proteins from the *Xanthomonas campestris* xanthan gum *gum* operon (GCA_001372255.1, contig B-1459_contig28 20,993-45,426), *Ensifer meliloti* succinoglycan *exo* operon (GCA_000006965.1, pSymB 1,155,981-1,190,869), and the *E. coli* colonic acid proteins (Wza, NP_416566.1; Wzc, NP_416564.4; Wzy, NP_310871.1; Wzz, NP_416531.4; Wzx, NP_310869.2) were used as a query for a blastp search. An additional blastp was done using the ezCAZy 2024 dataset as a custom database and using V0C-4 cluster proteins as a query.

For syntenic analyses (Fig. S3), the tool clinker (38) v0.0.32 was used using default settings for homology threshold (i.e., sequence identity of 30%). First, the *X. autotrophicus* V0C-4 gene cluster was used as a query against the entire genome sequence of the representative and type strains listed in Table S9 to locate where the homologous cluster was (i.e., where most of cluster genes colocalized within a region, singleton homologs not included). These strains were chosen as we had updated the genomes of these strains to long-read/hybrid assemblies, the majority of which formed a single complete contiguous chromosome. Most short-read only genomes that were available were too fragmented to be meaningfully included. This conserved gene cluster region from each genome with at least 3 homology hits was then extracted and all clusters were then re-analyzed using clinker for the final visualization in Fig. S3.

### Genetic Manipulations

The plasmid for expressing wild-type *X. autotrophicus* 7C *xsbA* was cloned via Golden Gate with BsaI by cloning from parts amplified from the XanthoMoClo plasmid toolkit(53). The *X. autotrophicus* 7C *xsbA* gene contains 2 internal BsaI sites, thus, Golden Gate junctions were designed at these sites, which when ligated, cure the BsaI site through a silent mutation that does not change the amino acid sequence of the protein, as described previously(53). Plasmid parts were amplified using TaKaRa GXL PrimeStar Polymerase (Cat No: R050B) according to the manufacturer’s instructions. PCR amplicons were purified by either New England Biolabs Monarch DNA Spin Gel Extraction Kit (Cat No: 1020L), New England Biolabs Monarch Spin PCR & DNA Cleanup Kit (Cat No: 1030L), or the Zymo Select-a-Size DNA Clean & Concentrator Kit (Cat No: D4080) by following the manufacturer instructions. The GenBank file for the final sequenced plasmid (long-read sequenced by Primordium, now Plasmidsaurus) is included in the Supplementary.

The *xsbA* homologs were seamlessly deleted from the *Xanthobacter* strains using homologous recombination, as described for *Pseudomonas*(54). The plasmids for knocking out *xsbA* homologs were cloned via Gibson by cloning homology arms upstream and downstream of the *xsbA* coding sequence into a plasmid backbone unable to replicate (pMB1 origin) but able to genomically integrate in the Xanthobacter strains via the homology. The SacB negative selection cassette was cloned from pJQ200SK(55), a gift from Michael Hynes (Addgene plasmid # 78497; http://n2t.net/addgene:78497; RRID:Addgene 78497). Plasmid parts were amplified and purified as described above. The final plasmids were assembled using new England Biolabs Gibson Assembly Master Mix (Cat No: E2611L) at 50°C for 1 hour following the manufacturer’s instructions. 1000 bp homology arms were used for the plasmids for *X. versatilis* Py2, *X. autotrophicus* 7C, *X. autotrophicus* V0C-4, and *X. tagetidis* TagT2C. For *X. flavus* GJ10, 500 bp homology arms were used instead due to a transposon insertion nearby its *xsbA* homolog. The GenBank files for these sequenced plasmids (long-read sequenced by Plasmidsaurus) are included in the Supplementary.

Plasmids were transformed into the strains using conjugation with either donor *E. coli* EPI300 ΔdapA pTA-Mob(56) or *E. coli* DH10B-MOB(53) (AddGene #229395). These donor strains were made chemically competent and transformed with the *xsbA* plasmid as described previously. Conjugations were done using saturated *Xanthobacter* cultures grown in 2xM1 (after 3 to 7 days of growth in 5 mL 2xM1 at 30°C) and overnight 1:100 diluted cultures of donor *E. coli* (grown at 37°C in 50 mL LB with 60 μg/mL diaminopimelic acid added (Dap60)). 100 μL of *Xanthobacter* culture and 50 or 100 μL of donor *E. coli* were plated on 2xM1 Dap60 plates. After incubation overnight at 30°C, cells were scraped with 1 mL of 2xM1 and replated onto 2xM1 plates containing either Kan30/200 (for plasmid expression of 7C *xsbA*) or 200 μg/mL spectinomycin (Spc200, for deletion plasmids). These plates were incubated for 7 days at 30°C, then colonies were picked and grown in liquid 2xM1 containing the appropriate antibiotic. For deletion constructs, cultures were passaged 1:100 into 5 mL 2xM1 without antibiotic and grown for 7 days at 30°C prior to plating a 10-fold serial dilution series on 2xM1 10% w/v sucrose (for negative selection via SacB). Slime-free colonies were visibly different but were also confirmed via whole-genome sequencing (long-read sequenced by Plasmidsaurus for *X. flavus* GJ10) or PCR using the conditions described above (*X. versatilis* Py2 – F: ATGGCGAAGATCGGCACCAG, R: CAGCGCTTTACCTACGTCTTCA; *X. autotrophicus* 7C – F: GAATGGTGTCGAGAAGATAGCC, R: CTTCACGTACGTCTTCACTACC; *X. autotrophicus* V0C-4 – F: CGAGCTTGTCGGCCAGGAACG, R: GTCTCGGTCCGGGTCAGATGG; *X. tagetidis* TagT2C – F: CGACCGGGATAGAGCGCGAG, R: CTCGGGCAAATGGTCGAGCACG)

### Transposon Libraries

Transposon libraries were generated using the pKMW7 barcoded Tn5 transposon library in conjugative donor *E. coli* (APA766)(57). An entire 1 mL glycerol stock of an OD_600_ greater than 1 was thawed and grown in 50 mL LB Dap60 Kan60 at 37°C. The saturated overnight culture was diluted 1:100 into a 500 mL culture of LB Dap60 Kan60 and harvested at an OD_600_ of around 0.3. The culture was spun down at 5,000 x *g* for 10 minutes, decanted, and resuspended to a final OD_600_ of 1. Saturated *X. autotrophicus* V0C-4 wild-type and Δ*xsbA* cultures were also spun down after being grown in 2xM1 for 5 days at 30°C. Both cultures were similarly spun down and resuspended to a final OD_600_ of 1.0. From these resuspensions, 100 μL of donor and recipient were plated onto 2xM1 Dap60 plates. These plates were grown overnight at 30°C, scraped with 1 mL of 2xM1, and 100 μL from the scraped volume was plated onto 2xM1 Kan30 8 times. In total, this was done 50 times in a single experiment for a total of approximately 400 final plates of *X. autotrophicus* V0C-4 wild-type or Δ*xsbA.* These plates were grown for 7 days at 30°C. Each plate was then scraped with 1 mL of 2xM1, pooled into a single flask, diluted in a new flask to an OD_600_ of 0.2 in 500 mL of 2xM1 Kan30, and grown overnight to a final OD_600_ of 1.2. A 2 mL volume of each pool was harvested for DNA isolation using the MasterPure Complete DNA and RNA Purification Kit from Biosearch Technologies (Cat No: MC85200) by following the manufacturer’s instructions under “Cell Samples – DNA Purificaiton Protocols for bacterial culture”.

The transposon library preparation protocol was followed as described for *Sphingobium*(58). To prepare the transposon library DNA for sequencing, isolated genomic DNA was diluted down to 25 ng/mL in 0.1x TE buffer. A 5 μL volume each of 100 μM Mod2_TS_Univ (ACGCTCTTCCGATC*T) and 100 μM Mod2_TruSeq (Phos-GATCGGAAGAGCACACGTCTGAACTCCAGTCA) were annealed to each other to form Y adapters using a PCR machine set to: 30 minutes at 37°C, ramp up to 97.5°C at 0.5°C/second, 97.5°C for 2 minutes and 30 seconds, ramp down to 20°C at 0.1°C/second, and hold at 4°C. This mix was then diluted to a concentration of 15 μM. 2.5 μL of the adapters were then ligated onto the gDNA using the NEBNext Ultra II FS DNA Library Prep (New England Biolabs Cat No: E6177S) according to manufacturer instructions for 300-450 bp. The eluted DNA off the size beads was then used in another PCR reaction that amplifies barcoded pieces, using the Nspacer_barseq_universal (ATGATACGGCGACCACCGAGATCTACACTCTTTCCCTACACGACGCTCTTCCGATCTNNNNNNGATGTCCACGAGGTCT) primer and the P7_Mod_TS_Index primer #2 (CAAGCAGAAGACGGCATACGAGATACATCGGTGACTGGAGTTCAGACGTGTGCTC TTCCGATCT) and #3 (CAAGCAGAAGACGGCATACGAGATGCCTAAGTGA CTGGAGTTCAGACGTGTGCTCTTCCGATCT). Using the Q5 High-Fidelity DNA polymerase (New England Biolabs Cat No: M0491L) according to manufacturer instructions, in a 100 μL reaction, 5 μL of each primer was added with 20 μL of eluted barcoded DNA and run with the following program: 98°C for 30 second; then 25 cycles of: 98°C for 10 seconds, 65°C for 30 seconds, and 72°C for 30 seconds; then a final step at 72°C for 2 minutes prior to holding at 4°C. The PCR reactions were cleaned up using the kits described above and eluted into 0.1x TE. The libraries were analyzed using Agilent D1000 ScreenTape, then diluted again down to 5 ng/μL, pooled, and sequenced using a NovaSeq X Plus (paired end sequenced, 2 x 150bp, 300 cycles).

For the transposon bioinformatic analyses, bcl2fastq was used to demultiplex reads. Reads were then processed using a set of custom scripts described below. We first analyzed the raw reads using a custom script identify_reads.c to extract the genomic region. Briefly, using the transposon model from the RB-TnSeq paper’s(57) implementation of the FEBA model (model_pKMW7_U), each read was first assessed for the correct transposon structure (N6-U1-N20-U2-IR-genomic DNA). Reads which matched this structure were then split into their constituent sequences. We aligned the genomic reads to the X. autotrophicus V0C4 reference genome using bowtie2 v2.5.4. For the slime-free strain, we used a reference genome lacking the xsbA gene coding sequence which we seamlessly deleted and confirmed via whole genome sequencing prior (via Plasmidsaurus Nanopore long-read whole genome shotgun sequencing). For every mapped read in the SAM alignment file, we use the mapped genome coordinates to identify if the read falls within a coding sequence (CDS) or in a non-coding region. For every read in a CDS, we further filter out reads that lie in the first and last 10% of the CDS body to only retain “central” insertions, which are more likely to have knocked out the gene(57). Due to a lack of barcode diversity (from the initial conjugation donor culture we used), instead of each uniquely barcoded read being counted as a unique insertion, we used unique insertion positions instead. While theoretically more than one unique transposon insertion could occur at the same position, it is unlikely, and filtering by unique insertion positions gave us a good measure of transposon insertion library diversity. The processed libraries corresponding to the wild-type and Δ*xsbA* (“VSF”) strains had different total numbers of unique central insertions (Supplementary Table 3) but widespread coverage in each (Fig. S4). For each library, we examined the distribution of the number of unique central insertions per gene (Fig. S5), which showed a sharp elbow in the plot. Using the kneed Python library, this elbow was determined to be at 10 insertions in each library. We interpret the elbow being that genes with low insertion counts (<10 insertions in V0C-4 and <10 insertions in VSF) have a lower than expected number of insertions, likely due to strong fitness effects from having a transposon insertion within the coding sequence of these genes. We filter out genes with <10 insertions in V0C-4 and <10 insertions in VSF for strain comparison downstream as we are interested not in essential genes across both strains, but genes with differential transposon insertions between the two strains. For functional annotations, we used the PGAP-provided RefSeq annotations (v6.11) for the *X. autotrophicus* V0C-4 genome, as well as Bakta v1.6.1 (59).

For calculating the fold-change in unqiue central insertions per gene, after the low insertion count genes were filtered, fold change was calculated as follows:

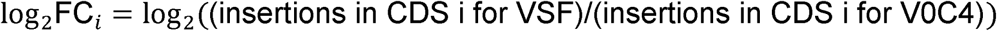

Since the total unique central insertion counts were not identical, we introduce a correction factor:

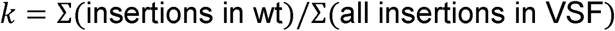

We then calculate the fold change in insertion counts as:

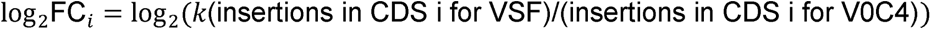

For calculating p-values, after correcting for library size and filtering low insertion count genes, we are left with 2248 genes with a sufficient number of insertions. We compute a p-value to test the null hypothesis that a gene’s log2FC value does not differ from the genome-wide background. We compute the Median Absolute Deviation (MAD) of the Log2(FC) across all genes that pass the filter. Assuming the Log2(FC) values are approximately normally distributed we use the appropriate scale factor (1.4826) to obtain a robust estimate of standard deviation. We compute a robust z-score for every CDS using the median of Log2(FC) computed across all genes that pass the filter:

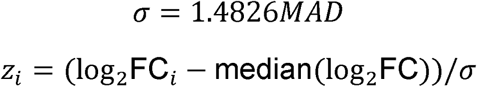

Finally, we compute a p-value for each filtered gene using the standard normal cumulative distribution function:

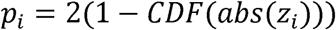

To control for the Family-wise Error Rate (FWER) at α=0.05, we compute a Bonferroni corrected significance threshold as 0.05/2248. 25 genes were identified as having significantly different insertion counts under the Bonferroni corrected p-values (Fig. 4A). Separately, we also applied the Benjamini-Hochberg False Discovery Rate (FDR) correction to the same set of p-values, by converting the p-values to q-values using the scipy.stats library false_discovery_control function (using the “bh” method). In total, 75 genes were identified as having significant insertions counts at FDR< 0.05 (Fig. S6).

The code implementing the overall pipeline can be found on GitHub (https://github.com/amoghpj/xanthobacter-transposon-insertion-analysis).

Individual clones that had aberrant slime production from the wild-type V0C-4 library were picked, restruck, and grown in 5 mL 2xM1 at 30°C for 7 days. Genomic DNA from these cultures was also extracted using the same method described above. Transposons (and their inserted genomic context) were captured by ligating them with an origin of replication, transforming into *E. coli* and sequenced as follows. The pMB1 origin from pXTK-Empty (from XanthoMoClo) was PCR amplified to add flanking NcoI cut sites using TaKaRa GXL by the conditions described above (F: GGTGGTCCATGGAGATCAAAGGATCTTCTTGAGATCCTTT, R: GGTGGTCCATGGCGCGTTGCTGGCGTTTTTCCAT). Genomic DNA from each transposon mutant and the PCR-amplified pMB1 origin were digested with NcoI-HF (New England Biolabs Cat No: R3193L), and ligated in a 1:1 ratio using T4 Ligase (New England Biolabs Cat No: M0202L) following manufacturer’s instructions. 25 μL ligated product was then transformed into chemically competent *E. coli* 5α (New England Biolabs Cat No: C2987U) according to manufacturer’s instructions. The entire recovered outgrowth was grown overnight in 5 mL LB Kan50 and miniprepped using the New England Biolabs Monarch Miniprep Kit (Cat No: T1010L/T1110L) according to manufacturer’s instructions. These minipreps were then sent for long-read plasmid sequencing to Plasmidsaurus for Nanopore sequencing. From the sequenced plasmids, 50 bp of flanking sequence either side of the Tn5 transposon was mapped to the *X. autotrophicus* V0C-4 reference genome in Geneious. Sequenced plasmids from the captured transposons are included in the Supplementary.

## Data Availability

The reads for *X. autotrophicus* 7CSF used for the mutation analysis can be found on NCBI at SRR40972798. The reads for the transposon libraries can be found on NCBI at SRR40981836 and SRR40981835. The sequences for the plasmids used in this study are included as GenBank files in the Supplementary.

## Acknowledgements

We would like to thank Dr. Bogumil Karas for the *E. coli* pTA-Mob strain and Dr. Valentine Trotter, Dr. Adam Arkin, and Dr. Adam Deutschbauer for the APA766 strain.

This work was supported by the Defense Advanced Research Projects Agency (DARPA), Award Number: N660012324014. The views, opinions, and/or findings expressed are those of the author and should not be interpreted as representing the official views or policies of the Department of Defense or the U.S. Government. This work was also supported in part by Kula Bio (A56981). M.P.M.S. was supported by a CGS-D/PGS-D from the Natural Sciences and Engineering Research Council of Canada. Funders had no role in study design, data collection and interpretation, or the decision to submit the work for publication.

No conflicts of interest are declared.

