## Supplementary material for "Identification and deletion of the extracellular polysaccharide slime pathway across the genus Xanthobacter": 2026_10_14__Supplementary.pdf

**Supplementary Table 1.** Mutations between *X. autotrophicus* 7C (GCF\_043194295.1) and *X. autotrophicus* 7CSF (GCF\_043194195.1). Mutations were determined by mapping 7CSF sequencing reads (SRR40972798) to the 7C genome using bowtie2 and identifying mutations above a quality score of 100 using bcftools mpileup, call, and filter functions. Mutations are relative to the 7C genome and sorted by quality. Initial mutation analysis was performed on the older RefSeq genomes at the time, prior to the release of the newer genomes now available (see Supplementary Table 9).

| Contig<br>(7C) | Position | Reference<br>(7C) | Mutation<br>(7CSF) | Type | Quality | Gene Affected |
| --- | --- | --- | --- | --- | --- | --- |
| NZ_JBAFVK010000016.1 | 24766 | C | A | Substitution | 225.417 | undecaprenyl-phosphate glucose phosphotransferase (V5F64_RS17230) |
| NZ_JBAFVK010000048.1 | 5 | GC | GCAGCGC | Indel | 132.172 | aldehyde dehydrogenase family protein (V5F64_RS22530) |
| NZ_JBAFVK010000053.1 | 5 | A | AATA | Indel | 130.328 | None |
| NZ_JBAFVK010000051.1 | 3 | TT | TTAATATT | Indel | 122.415 | None |
| NZ_JBAFVK010000053.1 | 3 | ATA | ATTATA | Indel | 121.376 | None |
| NZ_JBAFVK010000001.1 | 774294 | GCAAGG | GCAAGGGTGTCAAGG | Indel | 112.362 | None |
| NZ_JBAFVK010000041.1 | 6492 | CTATAT | CTATATAT | Indel | 111.139 | hypothetical protein (V5F64_RS22455) |
| NZ_JBAFVK010000016.1 | 113778 | GGCGCCC | GGCGCCCACCGTGCGCCC | Indel | 110.258 | None |
| NZ_JBAFVK010000052.1 | 6 | CTA | CTAGCACTA | Indel | 105.17 | None |
| NZ_JBAFVK010000043.1 | 3365 | GGCAT | GGCATGCAT | Indel | 102.303 | None |
| NZ_JBAFVK010000028.1 | 4 | AG | ATGGAG | Indel | 100.002 | None |

**Supplementary Figure 1.** Serial dilutions from cultures of diverse *Xanthobacter* strains across the genus plated on minimal media (modified Wiegel's minimal medium) with different carbon sources. Substrate concentrations are 0.5% v/v (for alcohols) or 0.5% w/v (for all others). Plates were grown for 7 days at 30°C. Serial dilutions were done in rich 2xM1 media to allow for enough residual nutrients to permit substrate metabolic switching. Species abbreviations are: XVE – *X. versatilis*, XOG – *X. oligotrophicus*, XAU – *X. autotrophicus*, XTO – *Xanthobacter* sp. T101, XNS – *X. nonsaccharivorans*, XLU – *X. lutulentifluminis*, XAM – *X. aminoxidans*, XFL – *X. flavus*, XWE – *X. wiegelii*, XCO – *X. cornucopiae*, XAG – *X. agilis*, XSE – *X. sediminis*, XAL – *X. albus*, XVA – *X. variabilis*, XPS – *X. pseudotagetidis*, XTA – *X. tagetidis*.

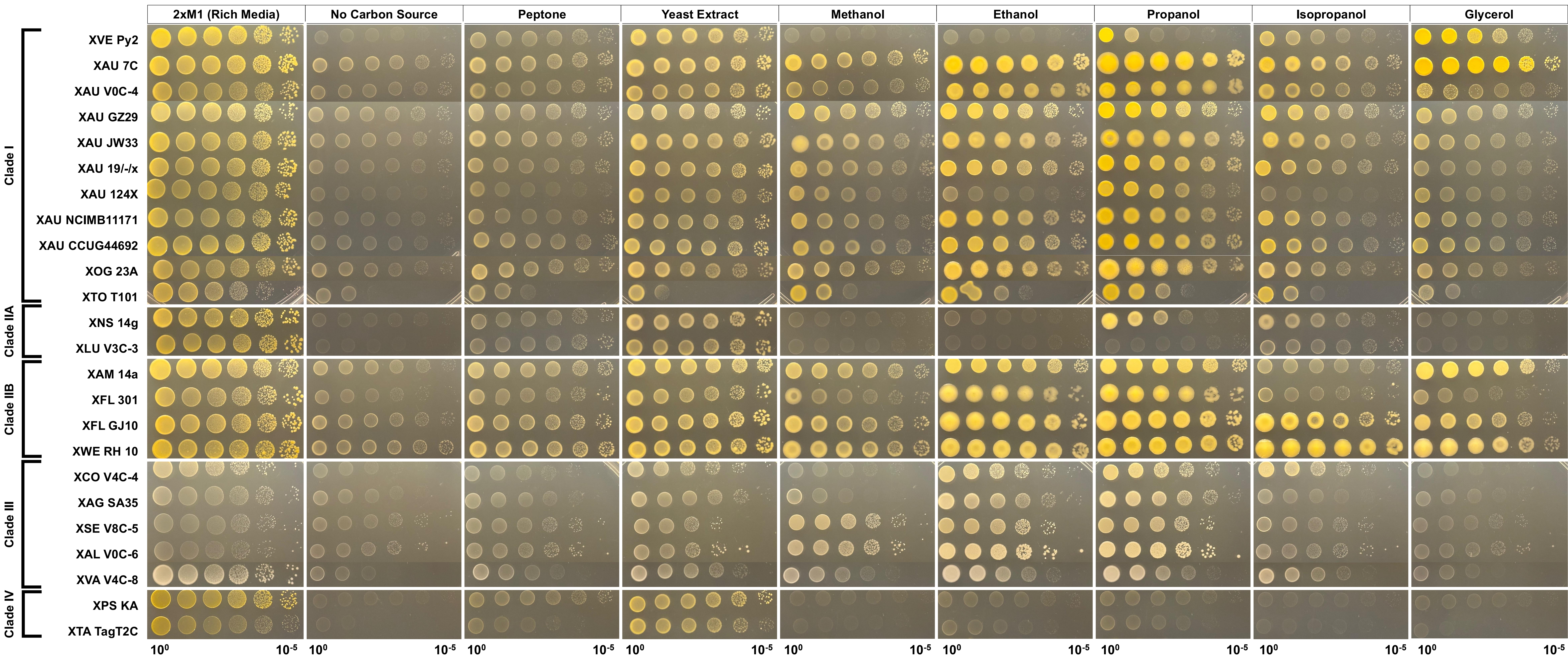

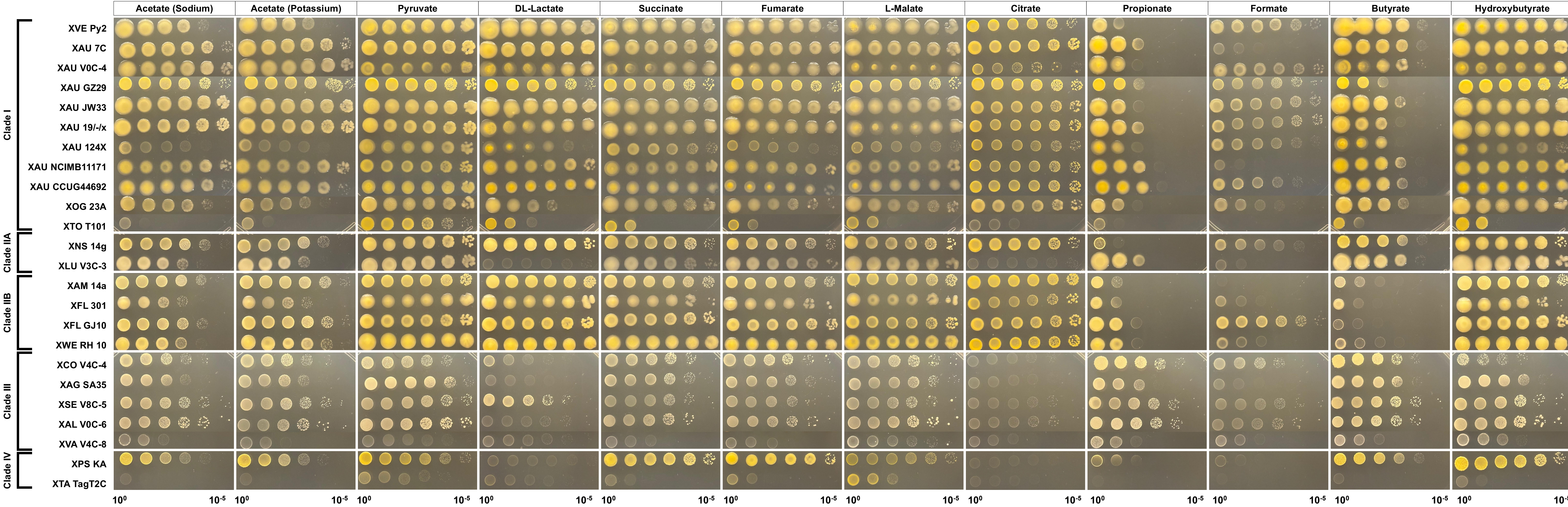

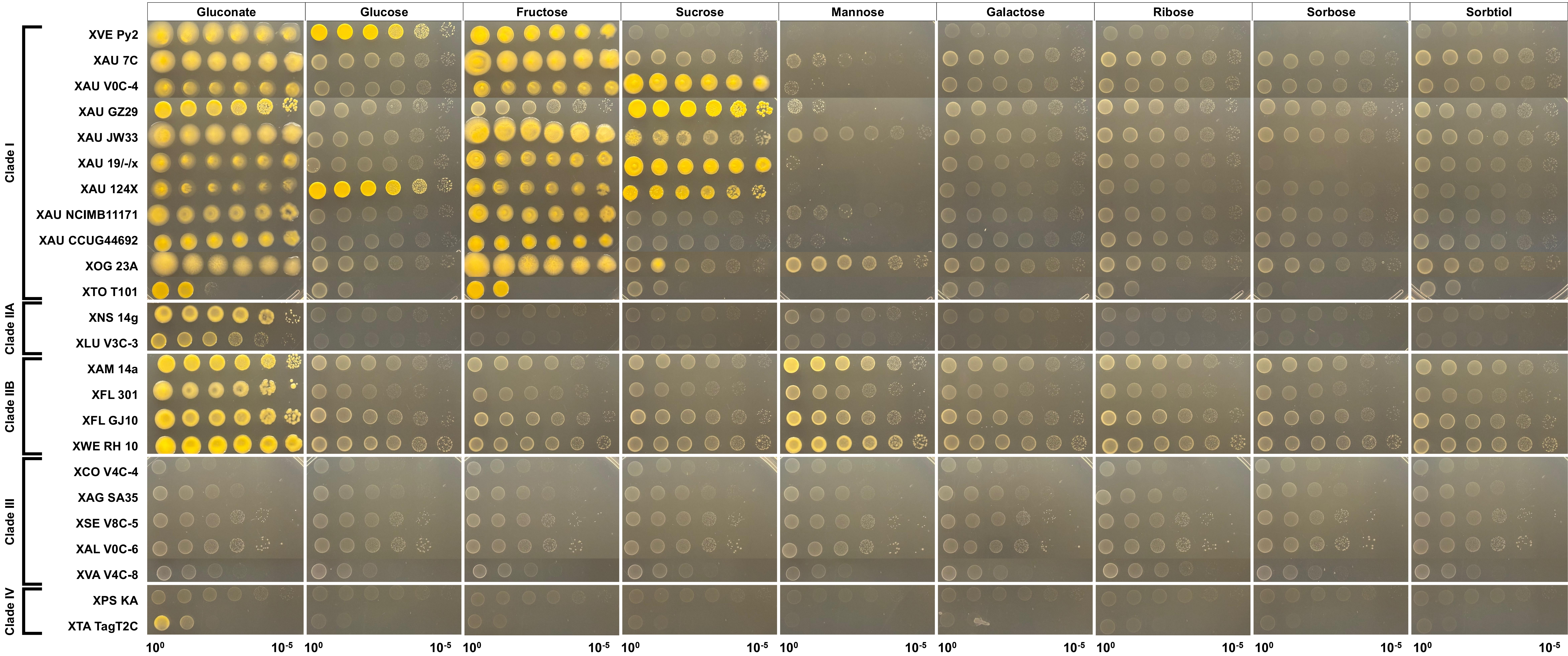

**Supplementary Table 2.** Presence of the exopolysaccharide biosynthetic pathway initiating protein undecaprenyl-phosphate glucose phosphotransferase *xsbA* across the genus *Xanthobacter*. Matches were identified using BLAST blastn (default settings) and blastp (default settings) in Geneious v2026.0. Query was *X. autotrophicus* 7C *xsbA* (NT: V5F64\_RS04325; AA: WP\_138398082.1) against custom BLAST nucleotide and protein databases from the genome sequences below. High identity *xsbA* homologs had an E-Value of 0 under both nucleotide and amino acid analyses with high query coverage (84-100% for nucleotide). Additional polyprenyl glycosylphosphotransferases are present in these genomes, however, they were clearly distinct from *xsbA* homologs as they did not have an E-Value of 0 to the query and had minimal query coverage (< ~20% nt) unlike high identity *xsbA* homologs.

| Species / Strain | Slime Induced / Produced | Clade | High Identity *xsbA* Hit NT (% Pairwise Identity) | High Identity *xsbA* Hit AA (% Pairwise Identity) | NCBI Genome |
| --- | --- | --- | --- | --- | --- |
| *X versatilis* Py2 | Yes | I | AC7W43_RS20380 (90.9%) | WP_506791320.1 (96.1%) | GCF_060823255.1 |
| *X oligotrophicus* 23A | Yes | I | V5F32_RS09470 (91.9%) | WP_188079529.1 (96.5%) | GCF_043194165.2 |
| *X autotrophicus* 7C | Yes | I | V5F64_RS04325 (100%) | WP_138398082.1 (100%) | GCF_043194295.2 |
| *X autotrophicus* V0C-4 | Yes | I | V5F34_RS19930 (99.5%) | WP_394075003.1 (99.6%) | GCF_043194495.2 |
| *Xanthobacter* sp. T101 | Minimal visible slimy change | I | V5F85_RS20705 (84.7%) | WP_394111566.1 (86.3%) | GCF_043194595.2 |
| *X nonsaccharivorans* 14g | Minimal visible slimy/waxy change | IIA | V5F59_RS13005 (77.5%) | WP_393978017.1 (73.4%) | GCF_043194755.2 |
| *X lutulentifluminis* V3C-3 | Yes | IIA | V5F49_RS19715 (81.8%) | WP_394114979.1 (81.5%) | GCF_043194675.2 |
| *X aminoxidans* 14a | Minimal visible waxy change | IIB | V5F57_RS10670 (76.2%) | WP_250165692.1 (72.6%) | GCF_043193655.2 |
| *X flavus* 301 | Slimy/waxy<br>visible<br>change | IIB | V5F82_RS19790<br>(75.5%) | WP_169124175.1<br>(72.0%) | GCF_043193365.2 |
| *X flavus* GJ10 | Slimy/waxy<br>visible<br>change | IIB | AC7W5B_RS10670<br>(74.3%) | WP_229643897.1<br>(71.3%) | GCF_060823235.1 |
| *X wiegelii* RH10 | Slimy/waxy<br>visible<br>change | IIB | V5F63_RS10050<br>(77.9%) | WP_394103047.1<br>(71.9%) | GCF_043193855.2 |
| *X cornucopiae* V4C-4 | No | III | No High Identity Homolog |  | GCF_043195065.2 |
| *X agilis* SA35 | No | III | No High Identity Homolog |  | GCF_043194985.2 |
| *X sediminis* V8C-5 | No | III | No High Identity Homolog |  | GCF_043195015.1 |
| *X albus* V0C-6 | No | III | No High Identity Homolog |  | GCF_043194895.2 |
| *X variabilis* V4C-8 | No | III | No High Identity Homolog |  | GCF_043831965.2 |
| *X pseudotagetidis* KA | Minimal<br>visible slimy<br>change | IV | V5F67_RS15805<br>(77.6%) | WP_394016851.1<br>(66.9%) | GCF_043194795.2 |
| *X tagetidis* TagT2C | Minimal<br>visible slimy<br>change | IV | V5F51_RS08465<br>(76.3%) | WP_170168514.1<br>(63.5%) | GCF_043194775.2 |

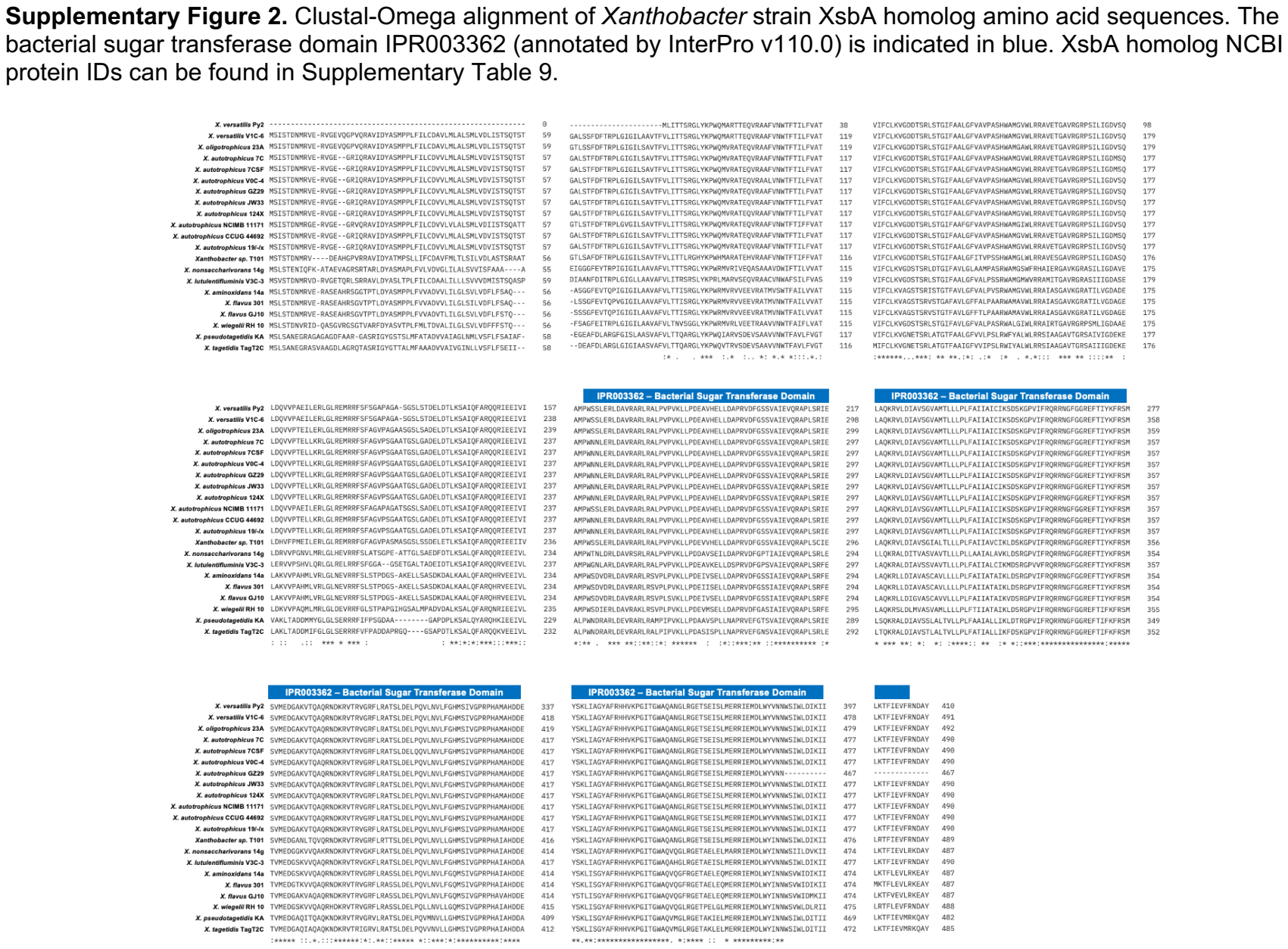

Supplementary Figure 2. Clustal-Omega alignment of Xanthobacter strain XsbA homolog amino acid sequences. The bacterial sugar transferase domain IPR003362 (annotated by InterPro v110.0) is indicated in blue. XsbA homolog NCBI protein IDs can be found in Supplementary Table 9.

**Supplementary Figure 3.** Gene conservation and syntenic structure conservation of the main slime biosynthetic gene cluster across slime-producing *Xanthobacter* strains analyzed by clinker using default settings. Genes are colored by group according to shared homology (i.e., homology indicated if sequence identity was above 30%). Homology group is indicated by color and number above or below the gene. Genes colored in dark green did not have any shared homologs across the strains but had annotated functions likely related to extracellular polysaccharide biosynthesis or were positioned adjacent to proteins that did. Genes colored in grey had annotated functions unrelated to extracellular polysaccharide biosynthesis. Genes colored in black were either transposases, recombinases, or integrases, (i.e., foreign element DNA).

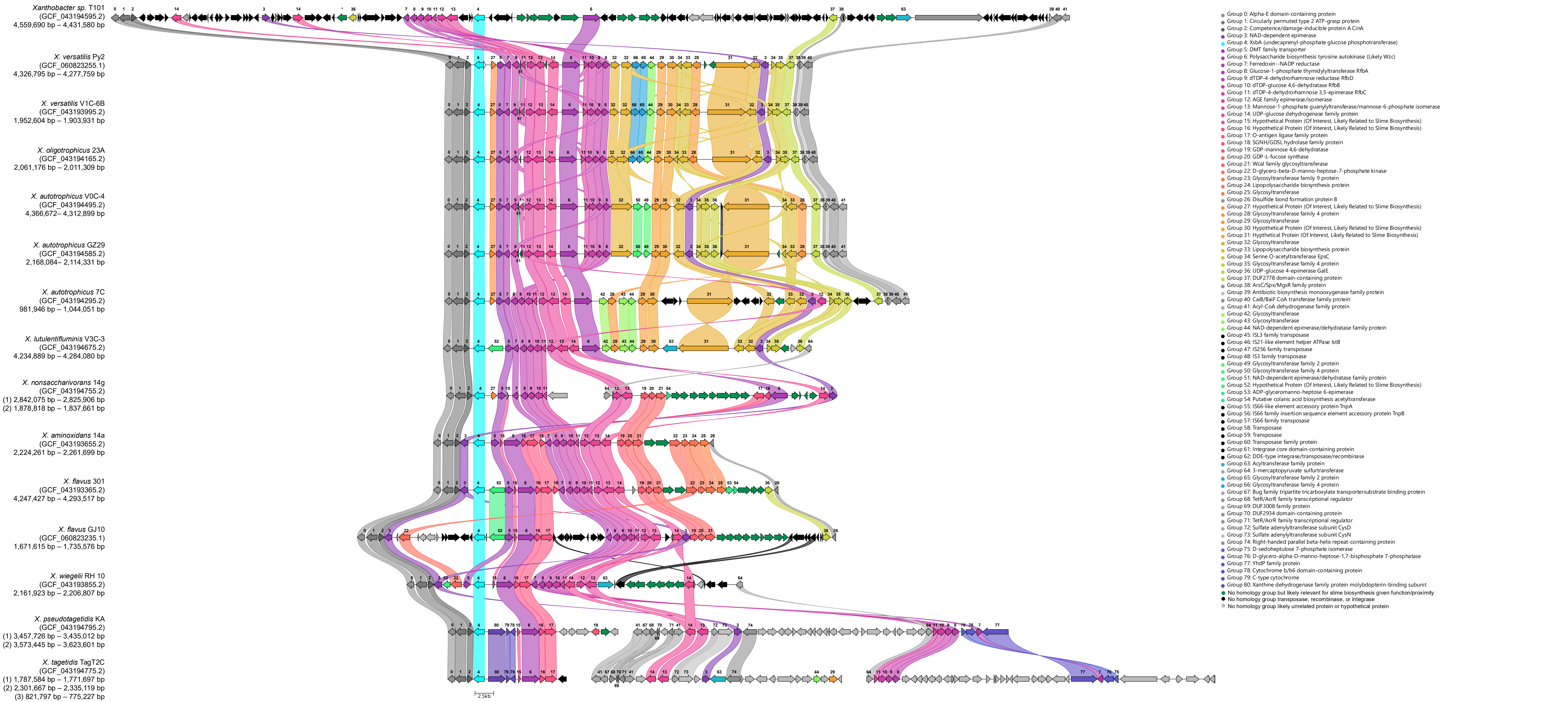

**Supplementary Table 3.** Tn5 transposon library statistics for *X. autotrophicus* V0C-4 and its slime-free knockout ($\Delta xsbA$, “VSF”).

| | *X. autotrophicus* V0C-4 | *X. autotrophicus* V0C-4 $\Delta xsbA$ (“VSF”) |
| --- | --- | --- |
| Tn5 reads successfully mapped to reference genome | 1,726,129 | 1,486,430 |
| Unique insertion sites (by position) | 87,520 | 145,402 |
| Unique insertion sites within coding sequence (CDS) | 73,456 | 124,766 |
| Unique central insertion sites (inner 10-90% of CDS) | 58,556 | 99,290 |
| Median unique central insertions per CDS | 4 | 6 |
| Mean unique central insertions per CDS | 11.46 | 19.44 |
| Max unique central insertions per CDS | 423 | 535 |

**Supplementary Table 4.** Transposon library insertion counts in *X. autotrophicus* V0C-4 $\Delta xsbA$ (“VSF”) and its wild-type parent. The table includes the RefSeq locus tag (“locus\_tag”), its annotated product (“refseq\_PGAP\_product”, “bakta\_product”), and unique insertions by position (“unique\_hit\_count\_V0C4/VSF”) and unique central insertions by position (insertions in the inner 10–90% of the coding sequence; “unique\_central\_hit\_count\_V0C4/VSF”).

**Supplementary Figure 4.** Transposon library insertion coverage across the genome in *X. autotrophicus* V0C-4 and its slime-free knockout ($\Delta xsbA$ “VSF”). The frequency of mapped transposon insertions is shown moving across the genome by chromosomal position (Mbp).

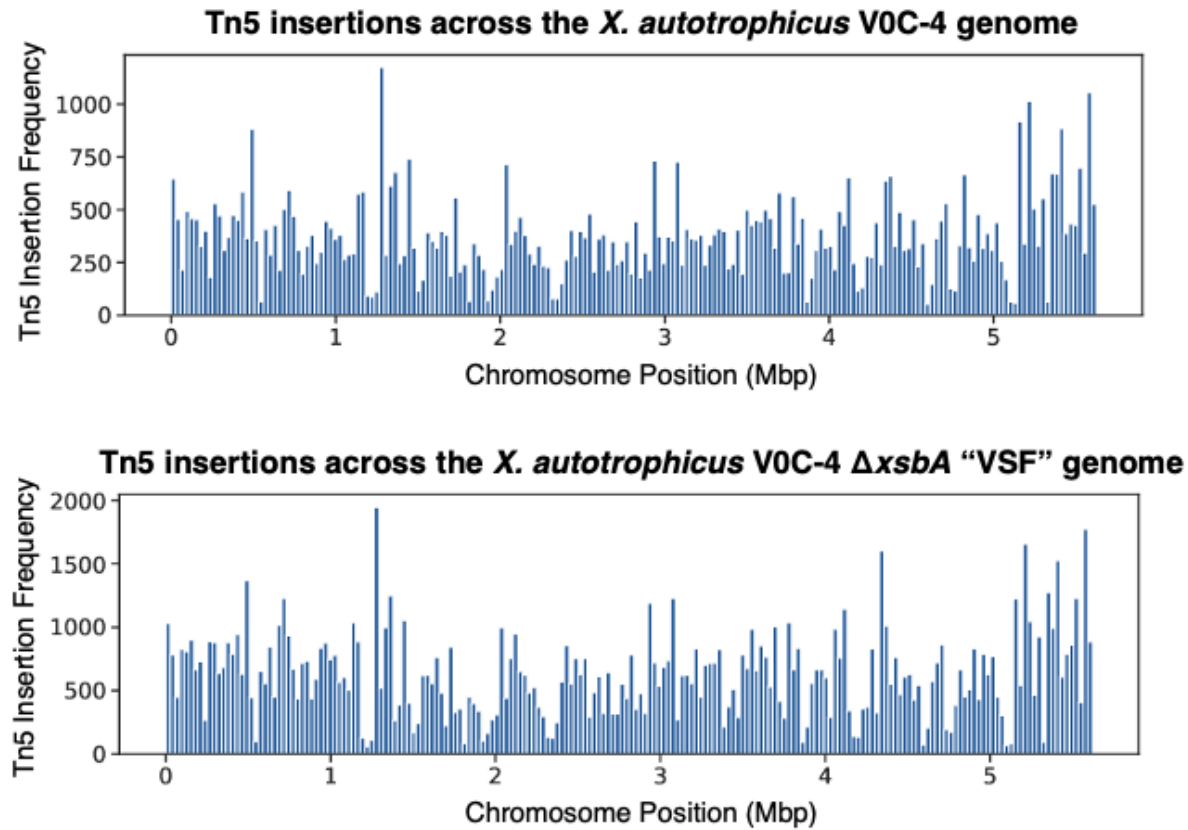

**Supplementary Figure 5.** Distribution of the number of unique central insertions per coding sequence in the transposon libraries for *X. autotrophicus* V0C-4 and its slime-free knockout ($\Delta xsbA$ “VSF”). The plots below show a sharp “elbow” at 10 insertions, as calculated using the kneed Python library. We interpret the elbow being that genes with low insertion counts (<10 insertions in V0C-4 and <10 insertions in VSF) have a lower than expected number of insertions, likely due to strong fitness effects from having an insertion within the coding sequence of these genes. We filter out genes with <10 insertions in V0C-4 and <10 insertions in VSF for strain comparison downstream as we are interested not in essential genes across both strains, but genes with differential transposon insertions between the two strains.

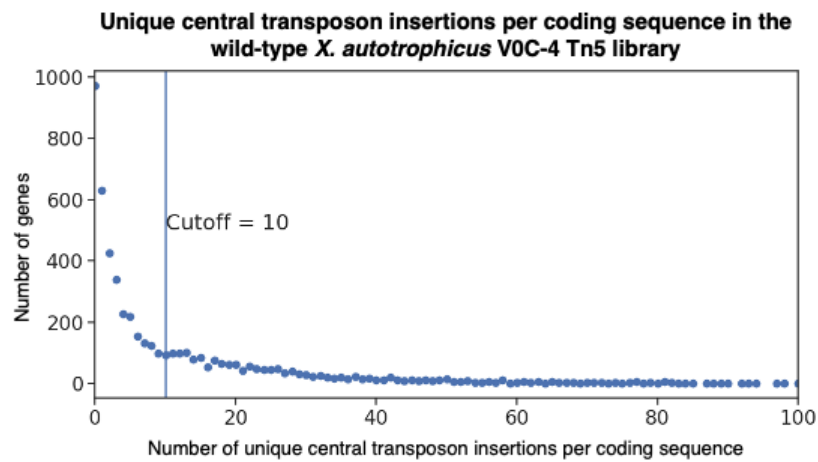

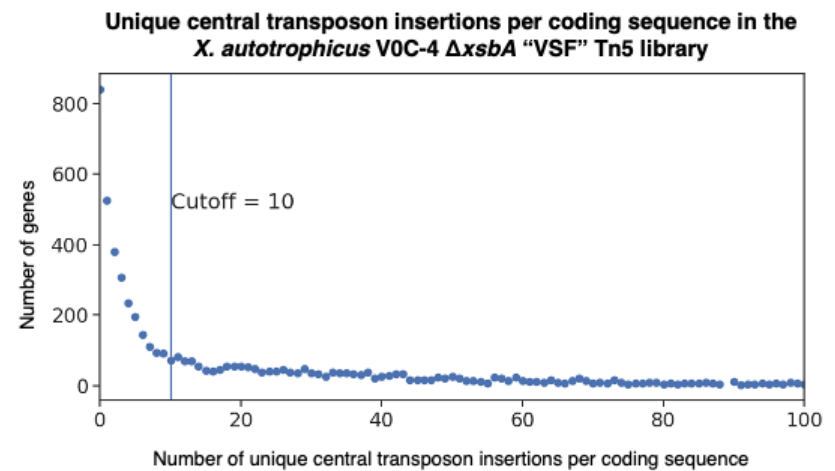

**Supplementary Table 5.** The log<sub>2</sub> fold change and calculated z-score and p/q-values of processed unique central transposon insertions per coding sequence in *X. autotrophicus* V0C-4 $\Delta xsbA$ “VSF” over its wild-type parent. Genes with less than <10 insertions in V0C-4 and <10 insertions in VSF are filtered out. The p-values were calculated using the standard normal cumulative distribution function with a Bonferroni corrected significance threshold (FWER). The q-values were calculated using the standard normal cumulative distribution function but corrected using the Benjamini-Hochberg False Discovery Rate (FDR). The columns include the gene RefSeq locus tag (“Locus tag”), the log<sub>2</sub> fold change (“Log<sub>2</sub>(VSF/V0C4)”), z score (“z”), Bonferroni corrected p-value (“=-LOG<sub>10</sub>(p)”), Benjamini-Hochberg False Discovery Rate corrected q-value (“=-LOG<sub>10</sub>(q)”), if the gene is considered significantly differentially inserted using Bonferroni corrected p-values (“is\_significant\_FWER”) or Benjamini-Hochberg FDR (“is\_significant\_FDR”), as well as the genes annotated product (by PGAP “refseq\_product” or Bakta “bakta\_product”).

**Supplementary Figure 6.** Log2 fold change of Tn5 transposon insertions per gene in the *X. autotrophicus* V0C-4 $\Delta xsbA$ strain library over wild-type. Genes that had significantly more insertions in one background than the other are colored in red ($p < 0.05$, Benjamini-Hochberg False Discovery Rate error corrected). The EPS slime biosynthetic pathway gene cluster is shaded in blue. The line at the top and bottom of the graph indicates genes with 0 insertions in one library, but 10 or more insertions in the other library.

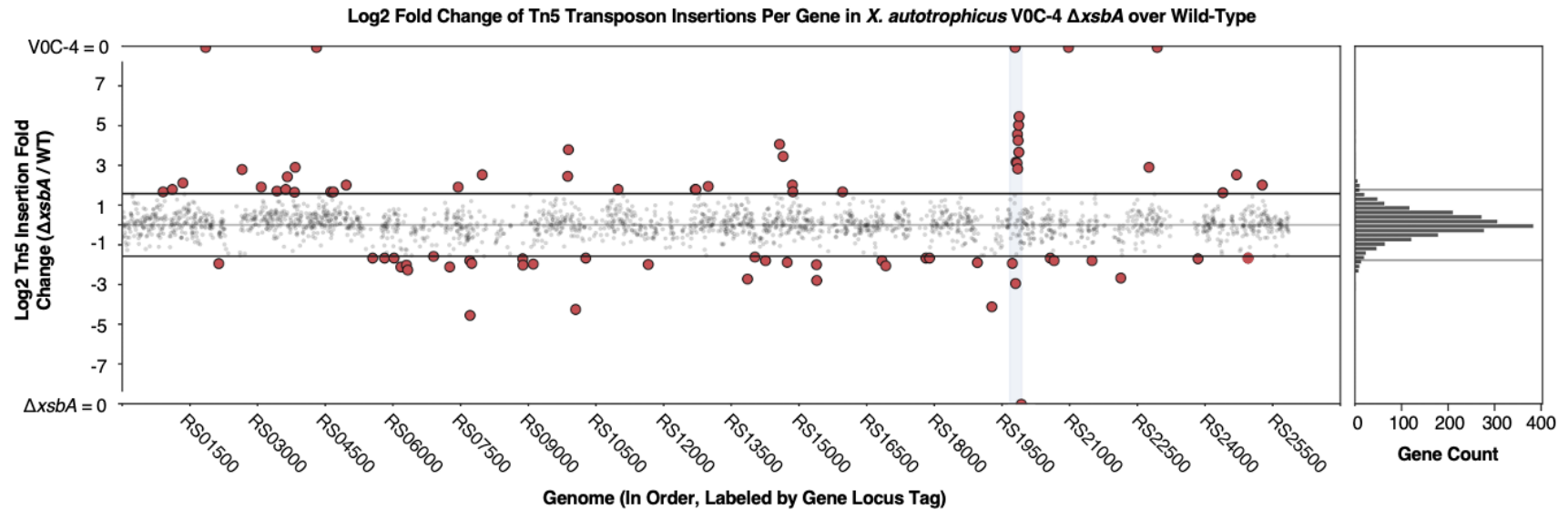

**Supplementary Table 6.** Genes with significantly more unique central transposon insertions in *X. autotrophicus* V0C-4 $\Delta xsbA$ “VSF” over wild-type. The gene p-values met the Bonferroni corrected significance threshold. Genes are indicated if they were found within the conserved 30 gene slime biosynthetic gene cluster (approximately 44 kbp).

| Locus | Gene Product (PGAP; Bakta) | In Slime Gene Cluster? | Log2 Insertion Fold-Change (VSF/V0C-4) | –Log10(p) |
| --- | --- | --- | --- | --- |
| V5F34_RS19880 | Polysaccharide biosynthesis tyrosine autokinase;<br>Lipopolysaccharide biosynthesis protein | Yes | 5.46 | 16 |
| V5F34_RS19870 | dTDP-glucose 4,6-dehydratase;<br>dTDP-glucose 4,6-dehydratase | Yes | 5.02 | 16 |
| V5F34_RS19845 | Glycosyltransferase family 2 protein;<br>Glycosyl transferase family 2 | Yes | 4.56 | 16 |
| V5F34_RS19860 | Glucose-1-phosphate thymidyltransferase RfbA;<br>Glucose-1-phosphate thymidyltransferase RfbA | Yes | 4.25 | 15.65 |
| V5F34_RS14570 | Polysaccharide biosynthesis/export family protein;<br>Periplasmic protein Wza involved in polysaccharide export, contains SLBB domain of the beta-grasp fold |  | 4.06 | 14.51 |
| V5F34_RS09890 | Hsp33 family molecular chaperone;<br>Hsp33 family molecular chaperone |  | 3.79 | 12.70 |
| V5F34_RS19875 | dTDP-4-dehydrorhamnose 3,5-epimerase;<br>dTDP-4-dehydrorhamnose 3,5-epimerase | Yes | 3.66 | 11.91 |
| V5F34_RS14650 | GNAT family N-acetyltransferase;<br>GNAT family N-acetyltransferase |  | 3.45 | 10.65 |
| V5F34_RS19810 | UDP-glucose 4-epimerase GalE;<br>UDP-glucose 4-epimerase GalE | Yes | 3.17 | 9.06 |
| V5F34_RS19835 | Hypothetical protein;<br>Hypothetical protein | Yes | 3.11 | 8.76 |
| V5F34_RS03835 | Cobaltochelatase subunit CobS;<br>Cobaltochelatase subunit CobS |  | 2.90 | 7.71 |
| V5F34_RS22760 | Hypothetical protein;<br>Hypothetical protein |  | 2.90 | 7.71 |
| V5F34_RS19850 | Glycosyltransferase family 4 protein;<br>Glycosyltransferase involved in cell wall<br>biosynthesis | Yes | 2.82 | 7.32 |
| V5F34_RS02655 | RrF2 family transcriptional regulator;<br>Rrf2 family transcriptional regulator |  | 2.79 | 7.16 |
| V5F34_RS07980 | TIGR03747 family integrating conjugative<br>element membrane protein;<br>Uncharacterized protein ORF SG67 |  | 2.52 | 5.97 |
| V5F34_RS24700 | ABC transporter substrate-binding protein;<br>Peripla-BP-6 domain-containing protein |  | 2.52 | 5.97 |
| V5F34_RS09875 | GcrA family cell cycle regulator;<br>GcrA cell cycle regulator |  | 2.45 | 5.65 |
| V5F34_RS03660 | Aminotransferase class I/II-fold pyridoxal<br>phosphate-dependent enzyme;<br>Aspartate/methionine/tyrosine<br>aminotransferase |  | 2.42 | 5.55 |

**Supplementary Table 7.** Genes with significantly more unique central transposon insertions in *X. autotrophicus* V0C-4 wild-type over $\Delta xsbA$ “VSF”. The gene p-values met the Bonferroni corrected significance threshold. Genes are indicated if they were found within the conserved 30 gene slime biosynthetic gene cluster (approximately 44 kbp).

| Locus | Gene Product (PGAP; Bakta) | In Slime Gene Cluster? | Log2 Insertion Fold-Change (V0C-4/VSF) | –Log10(p) |
| --- | --- | --- | --- | --- |
| V5F34_RS07710 | Glucose-6-phosphate dehydrogenase;<br>Glucose-6-phosphate dehydrogenase |  | 4.55 | 16 |
| V5F34_RS10050 | 6-phosphogluconolactonase;<br>6-phosphogluconolactonase/Glucosamine-6-phosphate isomerase/deaminase |  | 4.26 | 16 |
| V5F34_RS19275 | Hypothetical protein;<br>N/A |  | 4.12 | 15.35 |
| V5F34_RS19800 | Beta strand repeat-containing protein;<br>Hypothetical Protein | Yes | 2.95 | 8.26 |
| V5F34_RS15395 | ABC transporter permease;<br>ABC-type nitrate/sulfonate/bicarbonate transportsystem, permease component |  | 2.80 | 7.52 |
| V5F34_RS13860 | Methylmalonyl-CoA mutase subunit beta;<br>Hypothetical protein |  | 2.72 | 7.16 |
| V5F34_RS22135 | Oleate hydratase;<br>Oleate hydratase |  | 2.67 | 6.92 |

**Supplementary Table 8.** InterPro protein domains (v109.0) present within the proteins of the slime biosynthetic cluster in *X. autotrophicus* V0C-4. BLAST blastp similarity was also checked using *Xanthomonas campestris* xanthan gum cluster (GCA\_001372255.1, contig B-1459\_contig28 20,993-45,426), the *Ensifer/Sinorhizobium/Rhizobium meliloti* succinoglycan exo cluster (GCA\_000006965.1, pSymB 1,155,981-1,190,869), and Wza (NP\_416566.1), Wzc (NP\_416564.4), Wzx (NP\_310869.2), Wzy (NP\_310871.1), Wzz (NP\_416531.4), from *E. coli* colanic acid biosynthetic pathway. CAZy annotations were performed using the ezCAZy 2024 dataset as a custom BLAST database using the V0C-4 slime gene cluster as a query.

| Gene | Annotation (NCBI PGAP v6.11; Bakta v1.6.1) | Interpro Domains (v109.0) | BLAST Similarity blastp (wza/wzc/wzy/wzz/wzx, Xanthan gum Operon, Succinoglycan exo Operon) | CAZy BLAST Similarity blastp (ezCAZy 2024) |
| --- | --- | --- | --- | --- |
| V5F34_RS19930 (xsbA) | undecaprenyl-phosphate glucose phosphotransferase; undecaprenyl-phosphate glucose phosphotransferase | IPR003362, Bacterial sugar transferase (299-486); IPR017473, Undecaprenyl-phosphate glucose phosphotransferase, WcaJ (35-487); IPR017475, Exopolysaccharide biosynthesis polyprenyl glycosylphosphotransferase (35-488) | CEM57884.1 (gumD) *Xanthomonas campestris* (Pairwise Identity 46.3%; Query coverage: 56.08%; E-value $1.69 \times 10^{-78}$); CAC49469.1 (exoY) *Ensifer meliloti* (Pairwise Identity: 38.1%; Pairwise Identity: 38.1%; E-value $4.51 \times 10^{-34}$) | None |
| V5F34_RS19925 (ORF2) | hypothetical protein; hypothetical protein | None | None | None |
| V5F34_RS19920 (ORF3) | DMT family transporter; DMT family transporter rhaT | IPR037185, Multidrug transporter EmrE superfamily (40-142, 189-290); IPR000620, | None | None |
|  |  | EamA domain (6-138, 155-285) |  |  |
| V5F34_RS19915 (ORF4) | ferredoxin--NADP reductase; Flavodoxin/ferredoxin--NADP reductase fpr | IPR033892, Ferredoxin--NADP reductase, bacteria (1-254); IPR001709, Flavoprotein pyridine nucleotide cytochrome reductase (51-58, 111-130, 136-145, 215-223); | None | None |
| V5F34_RS19910 (ORF5) | rfbD dTDP-4-dehydrorhamnose reductase; dTDP-4-dehydrorhamnose reductase | IPR001433, Oxidoreductase FAD/NAD(P)-binding (112-229); IPR039261, Ferredoxin-NADP reductase (FNR), nucleotide-binding domain (100-253); IPR017927, FAD-binding domain, ferredoxin reductase-type (2-102); IPR017938, Riboflavin synthase-like beta-barrel (4-99); IPR008333, Flavoprotein pyridine nucleotide cytochrome reductase-like, FAD-binding domain (19-97) | CAC49465.1 (exoB) *Ensifer meliloti* (Pairwise Identity 23.8%; Query coverage: 43.77%; E-value $2.74 \times 10^{-02}$) | None |
| V5F34_RS19905 (ORF6) | NAD-dependent epimerase/dehydratase family protein; hypothetical protein | IPR036291, NAD(P)-binding domain superfamily (1-34); IPR001509, NAD-dependent | CEM57898.1 (NAD(P)H-dependent oxidoreductase) *Xanthomonas campestris* (Pairwise | None |
| | | epimerase/dehydratase [NAD dependent epimerase/dehydratase family] (3-37) | Identity 43.9%; Query coverage: 7.19%; E-value $2.40 \times 10^{-03}$; CAC49465.1 (exoB) *Ensifer meliloti* (Pairwise Identity 34.5%; Query coverage: 8.81%; E-value $4.92 \times 10^{-03}$) | |
| V5F34_RS19900 (ORF7) | dTDP-4-dehydrorhamnose 3,5-epimerase rfbC (frameshifted); hypothetical protein | IPR000888, dTDP-4-dehydrorhamnose 3,5-epimerase-like (6-71); IPR014710, RmlC-like jelly roll fold (5-72); IPR011051, RmlC-like cupin domain superfamily (6-71) | None | None |
| V5F34_RS19895 (ORF8) | AGE family epimerase/isomerase; Mannose-6-phosphate isomerase | IPR012341, Six-hairpin glycosidase-like superfamily (2-381); IPR010819, N-acylglucosamine 2-epimerase/Cellobiose 2-epimerase (35-373); IPR008928, Six-hairpin glycosidase superfamily (8-380) | None | GH13 (AAD00555.1; Pairwise Identity 21.9%; Query Coverage 43.64%; E-value $1.32 \times 10^{-02}$) |
| V5F34_RS19890 (ORF9) | mannose-1-phosphate guanylyltransferase/mannose -6-phosphate isomerase; mannose-1-phosphate guanylyltransferase/mannose -6-phosphate isomerase cpsB | IPR029044, Nucleotide-diphospho-sugar transferases (16-294); IPR001538, Mannose-6-phosphate isomerase, type II, C-terminal (364- | None | None |
|  |  | 478); IPR006375, Mannose-1-phosphate guanylyltransferase/mannose-6-phosphate isomerase (16-482); IPR011051, RmlC-like cupin domain superfamily (202-483); IPR005835, Nucleotidyl transferase domain (18-299); IPR051161, Mannose-6-phosphate isomerase type 2 (15-477); IPR049577, GDP-mannose pyrophosphorylase, N-terminal domain (16-291); IPR014710, RmlC-like jelly roll fold (361-466); IPR054566, MannoseP isomerase/GMP-like, beta-helix domain (310-360) |  |  |
| V5F34_RS19885 (ORF10) | UDP-glucose dehydrogenase family protein; UDP-glucose 6-dehydrogenase ugd | IPR036220, UDP-glucose/GDP-mannose dehydrogenase, C-terminal domain superfamily (302-428); IPR014027, UDP-glucose/GDP-mannose dehydrogenase, C-terminal (317-418); | None | None |
|  |  | IPR028357, UDP-glucose 6-dehydrogenase, bacterial type (1-434); IPR008927, 6-phosphogluconate dehydrogenase-like, C-terminal domain superfamily (201-294); IPR014026, UDP-glucose/GDP-mannose dehydrogenase, dimerization (201-293); IPR017476, UDP-glucose/GDP-mannose dehydrogenase (1-414); IPR001732, UDP-glucose/GDP-mannose dehydrogenase, N-terminal (1-188); IPR036291, NAD(P)-binding domain superfamily (1-188) |  |  |
| V5F34_RS19880 (ORF11) | polysaccharide biosynthesis tyrosine autokinase; Lipopolysaccharide biosynthesis protein | IPR027417, P-loop containing nucleoside triphosphate hydrolase (477-750); IPR005702, Tyrosine-protein kinase Wzc-like, C-terminal domain (559-740); IPR003856, Polysaccharide chain length determinant, N-terminal domain [Wzz] | NP_416564.4 (wzc) *E. coli* (Pairwise Identity 23.2%; Query Coverage 49.17%; E-value $3.92 \times 10^{-19}$); NP_416531.4 (wzz) *E. coli* (Pairwise Identity 29.0%; Query coverage: 21.17%; E-value $6.15 \times 10^{-03}$); CAC49486.1 (exoP) | None |
| | | (27-118); IPR032807, Tyrosine-protein kinase, G-rich domain (407-477); IPR050445, Bacterial polysaccharide biosynthesis and export (21-740); IPR025669, AAA domain (565-686) | *Ensifer meliloti* (Pairwise Identity: 36.7%; Query coverage: 96.06%; E-value $2.29 \times 10^{-144}$) | |
| V5F34_RS19875 (ORF12) | rfbC dTDP-4-dehydrorhamnose 3,5-epimerase; dTDP-4-dehydrorhamnose 3,5-epimerase | IPR011051, RmlC-like cupin domain superfamily (1-183); IPR000888, dTDP-4-dehydrorhamnose 3,5-epimerase-like (3-178); IPR014710, RmlC-like jelly roll fold (1-184) | None | None |
| V5F34_RS19870 (ORF13) | rfbB dTDP-glucose 4,6-dehydratase; dTDP-glucose 4,6-dehydratase rfbB | IPR016040, NAD(P)-binding domain (4-320); IPR005888, dTDP-glucose 4,6-dehydratase (1-330); IPR036291, NAD(P)-binding domain superfamily (1-340) | CEM57898.1 (NAD(P)H-dependent oxidoreductase)<br>*Xanthomonas campestris* (Pairwise Identity 24.1%; Query coverage: 73.44%; E-value $1.43 \times 10^{-04}$);<br>CAC49465.1 (exoB)<br>*Ensifer meliloti* (Pairwise Identity 28.0%; Query coverage: 97.57%; E-value $4.51 \times 10^{-19}$);<br>CAC49476.1 (exol)<br>*Ensifer meliloti* (Pairwise Identity: | None |
| | | | 31.7%; Pairwise Identity: 31.7%; E-value $1.28 \times 10^{-02}$) | |
| V5F34_RS19865 (ORF14) | rfbD dTDP-4-dehydrorhamnose reductase; dTDP-4-dehydrorhamnose reductase | IPR029903, RmlD-like, substrate binding domain (1-290); IPR005913, dTDP-4-dehydrorhamnose reductase (2-291); IPR036291, NAD(P)-binding domain superfamily (1-291) | CEM57898.1 (NAD(P)H-dependent oxidoreductase) *Xanthomonas campestris* (Pairwise Identity 30.6%; Query coverage: 19.38%; E-value $2.64 \times 10^{-03}$) | None |
| V5F34_RS19860 (ORF15) | glucose-1-phosphate thymidyltransferase RfbA; glucose-1-phosphate thymidyltransferase RfbA | IPR005907, Glucose-1-phosphate thymidyltransferase, short form (1-240); IPR029044, Nucleotide-diphospho-sugar transferases [Spore Coat Polysaccharide Biosynthesis Protein SpsA; Chain A] (1-286); IPR005835, Nucleotidyl transferase domain (2-238) | CAC49485.1 (exoN) *Ensifer meliloti* (Pairwise Identity: 27.5%; Query coverage: 75.50%; E-value $8.85 \times 10^{-16}$) | None |
| V5F34_RS19855 (ORF16) | glycosyltransferase; hypothetical protein | IPR017853, Glycoside hydrolase superfamily (486-818) | None | 56 hits, top hit: GT4 (APK80348.1; Pairwise Identity 31.7%; Query Coverage 14.62%; E- |
| | | | | value $1.83 \times 10^{-11}$) |
| V5F34_RS19850 (ORF17) | glycosyltransferase family 4 protein; Glycosyltransferase involved in cell wall biosynthesis | IPR050194, Glycosyltransferase group 1 (193-359) | CEM57889.1 (gumI) *Xanthomonas campestris* (Pairwise Identity 23.4%; Query coverage: 45.71%; E-value $1.49 \times 10^{-03}$) | 34 hits, top hit: GT4 (NP_595519.1; Pairwise Identity 25.6%; Query Coverage 51.89%; E-value $3.49 \times 10^{-07}$) |
| V5F34_RS19845 (ORF18) | glycosyltransferase family 2 protein; Glycosyl transferase family 2 | IPR029044, Nucleotide-diphospho-sugar transferases [Spore Coat Polysaccharide Biosynthesis Protein SpsA; Chain A] (4-263) | CAC49482.1 (exoA) *Ensifer meliloti* (Pairwise Identity: 24.0%; Query coverage: 54.98%; E-value $2.15 \times 10^{-03}$); CAC49484.1 (exoO) *Ensifer meliloti* (Pairwise Identity: 29.6%; Query coverage: 37.54%; E-value $2.49 \times 10^{-10}$) | 44 hits, top hit: GT2 (AAK47706.1; Pairwise Identity 28.9%; Query Coverage 80.00%; E-value $1.40 \times 10^{-24}$) |
| V5F34_RS19840 (ORF19) | Glycosyltransferase; Glycos-transf-1 domain-containing protein | IPR001296, Glycosyl transferase, family 1 (187-328); IPR028098, Glycosyltransferase subfamily 4-like, N-terminal domain (15-180); IPR050194, Glycosyltransferase group 1 (6-364) | CEM57888.1 (gumH) *Xanthomonas campestris* (Pairwise Identity 25.3%; Query coverage: 37.53%; E-value $7.57 \times 10^{-04}$) | 100+ hits, top hit: GT4 (BAC09142.1; Pairwise Identity 33.3%; Query Coverage 45.45%; E- |
| | | | | value $1.68 \times 10^{-19}$) |
| V5F34_RS19835 (ORF20) | hypothetical protein; hypothetical protein | None | None | None |
| V5F34_RS19830 (ORF21) | cellulase family glycosylhydrolase; Putative glycosyl hydrolase | IPR017853, Glycoside hydrolase superfamily (50-366); IPR001547, Glycoside hydrolase, family 5 (158-311); IPR051923, Glycosyl Hydrolase Family 39 (41-367) | None | 7 weak hits, top hit: GH5 (ABI94085.1; Pairwise Identity 27.3%; Query Coverage 22.93%; E-value $1.34 \times 10^{-03}$) |
| V5F34_RS19825 (ORF22) | NAD-dependent epimerase; Nucleoside-diphosphate-sugar epimerase wcaG | IPR036291, NAD(P)-binding domain superfamily (1-334); IPR001509, NAD-dependent epimerase/dehydratase (4-234) | CEM57898.1 (NAD(P)H-dependent oxidoreductase) *Xanthomonas campestris* (Pairwise Identity 23.8%; Query coverage: 98.75%; E-value $6.95 \times 10^{-09}$); CAC49465.1 (exoB) *Ensifer meliloti* (Pairwise Identity 22.1%; Query coverage: 96.05%; E-value $1.37 \times 10^{-21}$) | None |
| V5F34_RS19820 (ORF23) | serine O-acetyltransferase EpsC; Serine acetyltransferase | IPR053376, Serine acetyltransferase (34-182); IPR001451, Hexapeptide repeat (130-163); IPR045304, | None | None |
|  |  | Serine acetyltransferase, LbH domain (79-177); IPR011004, Trimeric LpxA-like superfamily (39-181) |  |  |
| V5F34_RS19815 (ORF24) | glycosyltransferase family 4 protein; Glycosyltransferase | IPR001296, Glycosyl transferase, family 1 (224-365); IPR028098, Glycosyltransferase subfamily 4-like, N-terminal domain (19-211) | None | 97 hits, top hit: GT4 (BAC09143.1; Pairwise Identity 24.4%; Query Coverage 95.45%; E-value $4.67 \times 10^{-13}$) |
| V5F34_RS19810 (ORF25) | UDP-glucose 4-epimerase GalE; UDP-glucose 4-epimerase GalE | IPR005886, UDP-glucose 4-epimerase (3-321); IPR036291, NAD(P)-binding domain superfamily (1-322); IPR001509, NAD-dependent epimerase/dehydratase (3-252) | CEM57898.1 (NAD(P)H-dependent oxidoreductase) *Xanthomonas campestris* (Pairwise Identity 21.9%; Query coverage: 44.06%; E-value $3.65 \times 10^{-04}$); CAC49465.1 (exoB) *Ensifer meliloti* (Pairwise Identity 44.2%; Query coverage: 97.87%; E-value $5.35 \times 10^{-85}$) | None |
| V5F34_RS19805 (ORF26) | IS3 family transposase; Not annotated | None | None | None |
| V5F34_RS19800 (ORF27) | beta strand repeat-containing protein; hypothetical protein | IPR017853, Glycoside hydrolase superfamily | None | None |
| | | (25-289); Additional domains for the catalytic TIM $\beta/\alpha$ barrel (498-613, 614-723, 724-817, 845-965, 966-1084, 1085-1199, 1200-1321, 1322-1433, 1434-1556, 1627-1749, 1753-1859) | | |
| V5F34_RS19795 (ORF28) | serine O-acetyltransferase; Serine acetyltransferase | IPR011004, Trimeric LpxA-like superfamily (15-167); IPR005881, Serine O-acetyltransferase (3-176) | None | None |
| V5F34_RS19790 (ORF29) | lipopolysaccharide biosynthesis protein; Polysaccharide biosynthesis protein | IPR050833, Polysaccharide Biosynthesis and Transport (16-406) [PST FAMILY PREDICTED COLANIC ACID TRANSPORTER] | None | None |
| V5F34_RS19785 (ORF30) | glycosyltransferase family 4 protein; Glycos-transf-1 domain-containing protein | IPR001296, Glycosyl transferase, family 1 (191-324); IPR028098, Glycosyltransferase subfamily 4-like, N-terminal domain (107-173) | CEM57888.1 (gumH) *Xanthomonas campestris* (Pairwise Identity 26.9%; Query coverage: 52.76%; E-value $1.37 \times 10^{-07}$) | 75 hits, top hit: GT4 (AGH30276.1; Pairwise Identity 37.0%; Query Coverage 31.64%; E-value $3.91 \times 10^{-14}$) |

**Supplementary Figure 7.** Culture pellets from wild-type and $\Delta xsbA$ strains of diverse slime-producing *Xanthobacter*, including *X. versatilis* Py2, *X. autotrophicus* 7C, *X. autotrophicus* V0C-4, *X. flavus* GJ10, and *X. tagetidis* TagT2C. Cultures were grown in either 2xM1 or R2A rich media for 7 days prior to centrifugation at 5,000 x g for 20 minutes.

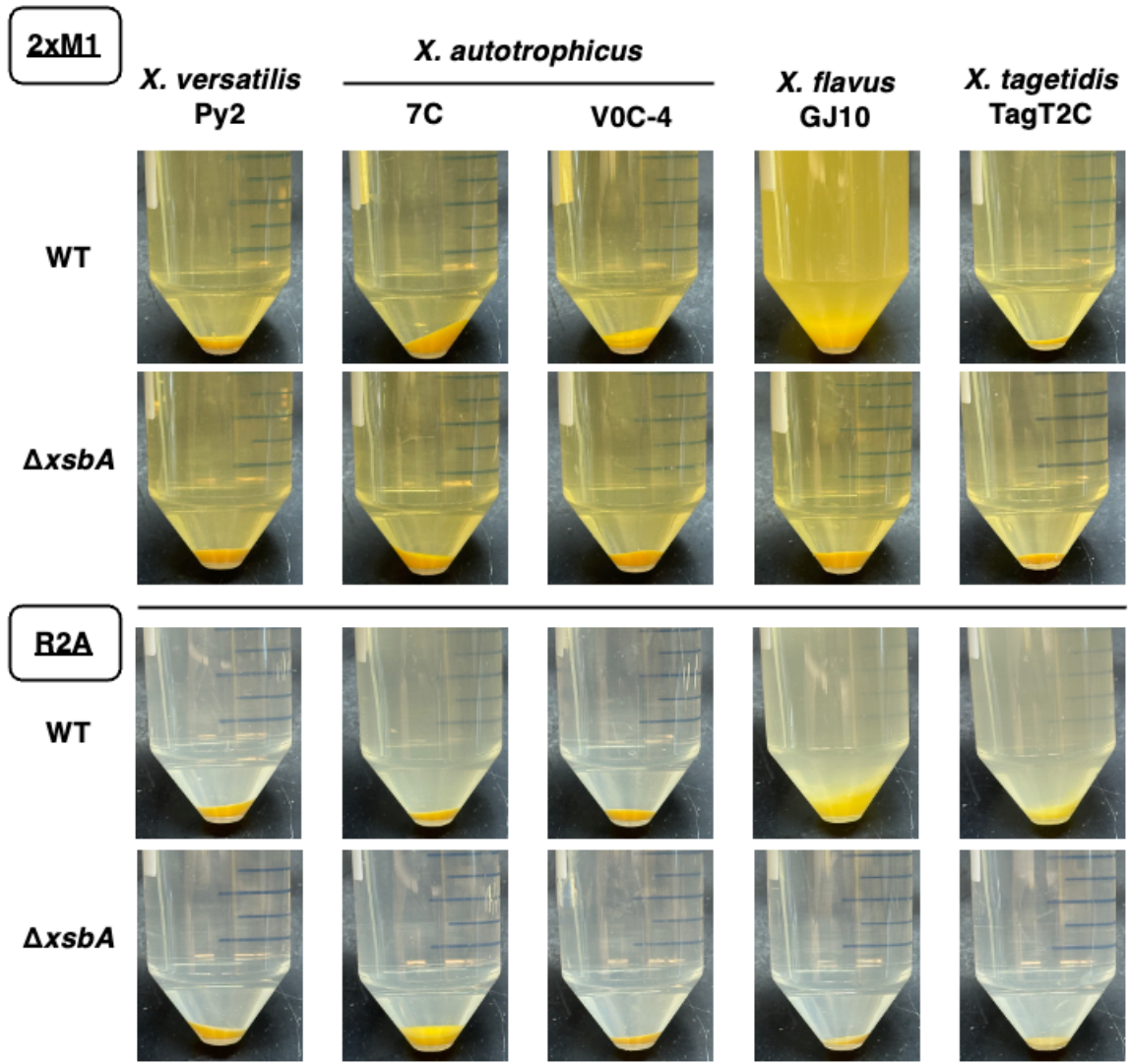

**Supplementary Figure 8.** Optical density (OD<sub>600</sub>) measurements from liquid cultures of *Xanthobacter* wild-type and $\Delta xsbA$ strains grown in inorganic minimal media with an autotrophic gas mixture bubbled in (~2% O<sub>2</sub>, ~10% CO<sub>2</sub>, ~3% H<sub>2</sub>, rest N<sub>2</sub>). In this condition, strains must grow fully autotrophically and fix nitrogen.

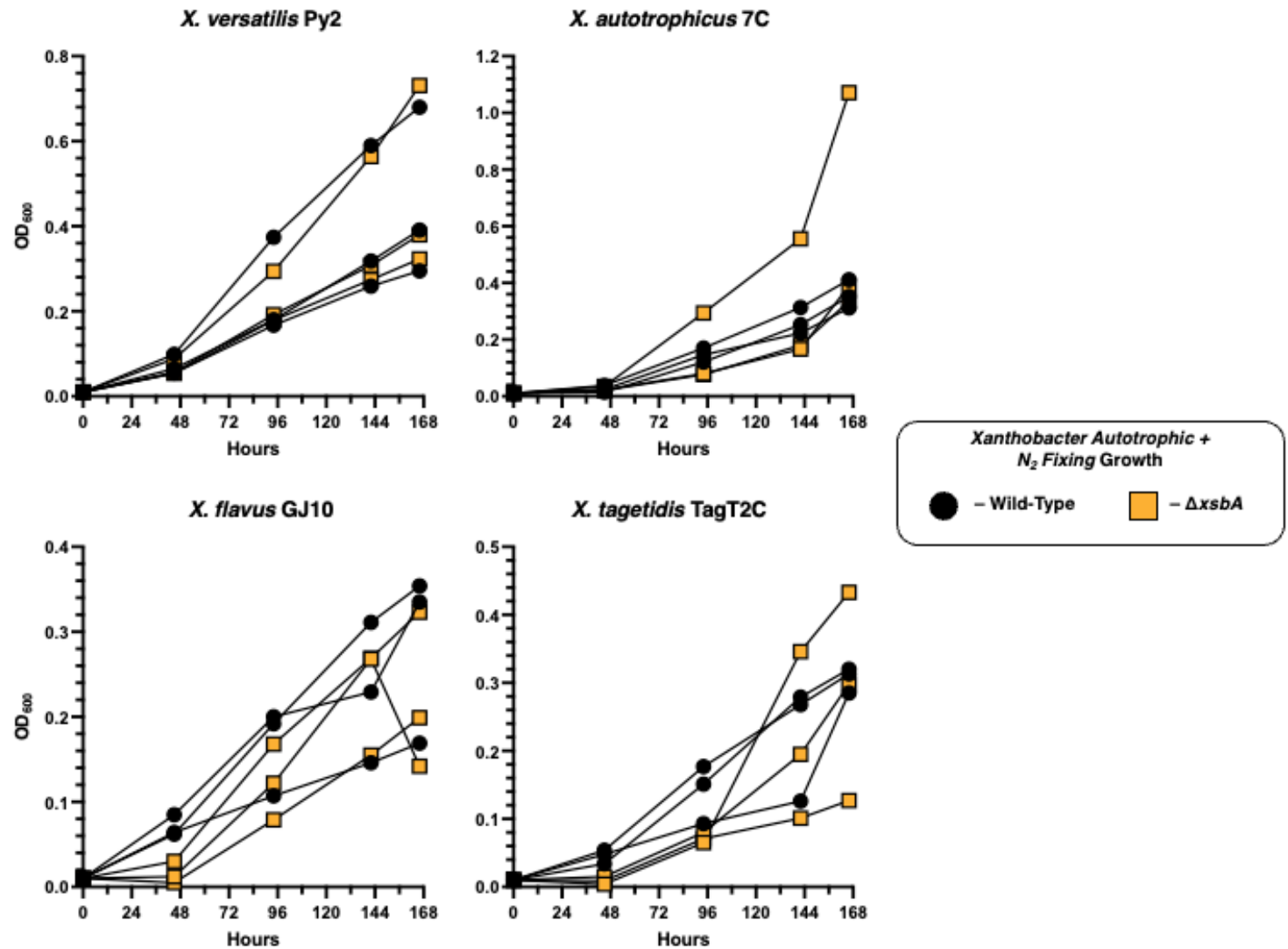

**Supplementary Figure 9.** Serial dilutions of wild-type and $\Delta xsbA$ *Xanthobacter* strains spot plated onto either inorganic minimal media (with or without bicarbonate), modified Wiegel's minimal media (with or without bicarbonate) or 2xM1 rich media. The presence or absence of nitrogen (3 g/L ammonium sulfate) or carbon (0.5% w/v succinate) is indicated. Plates were grown for 7 days at 30 °C either under ambient air or within a sealed bag continuously pumped with an autotrophic gas mixture (~2% O<sub>2</sub>, ~10% CO<sub>2</sub>, ~3% H<sub>2</sub>, rest N<sub>2</sub>). Without nitrogen added to the plate, strains must fix nitrogen. Without carbon added to the plate, strains must fix CO<sub>2</sub> (autotrophic growth).

IMM – Bicarbonate

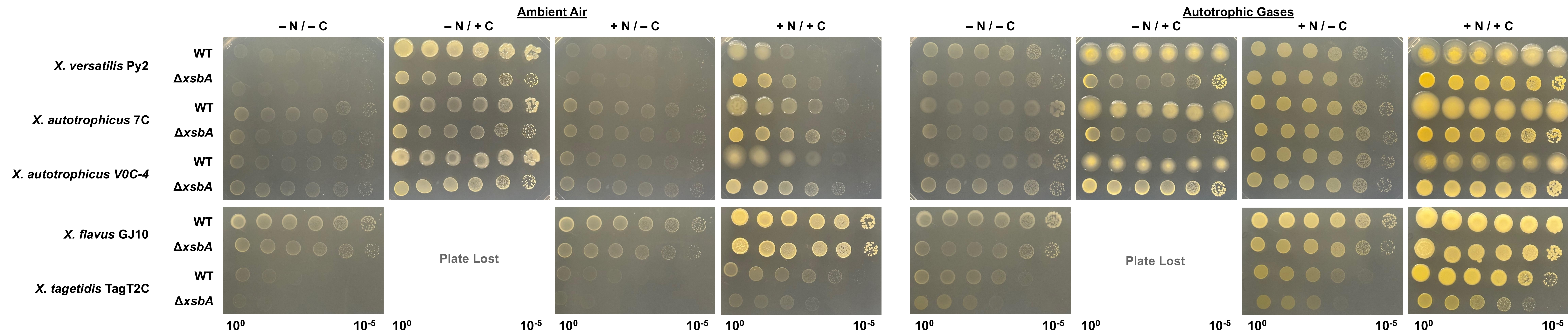

IMM + Bicarbonate

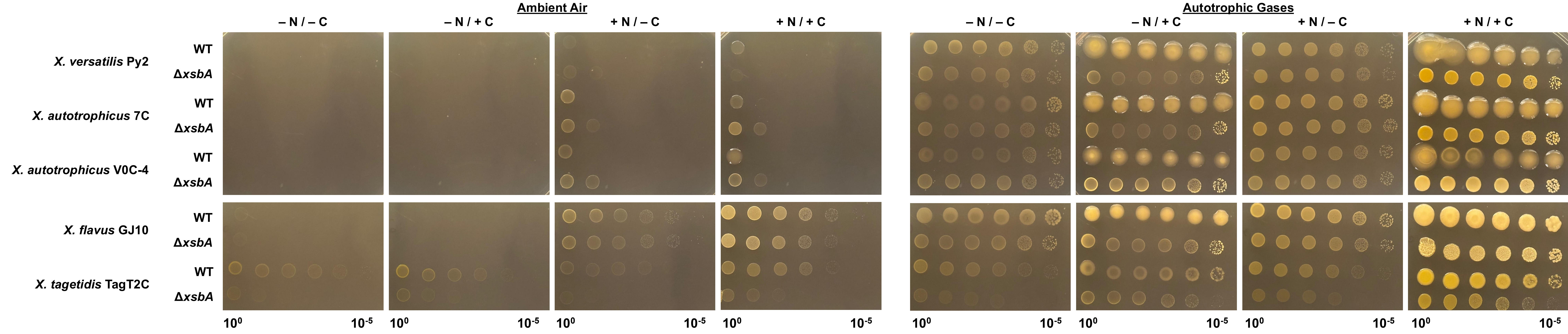

WMM – Bicarbonate

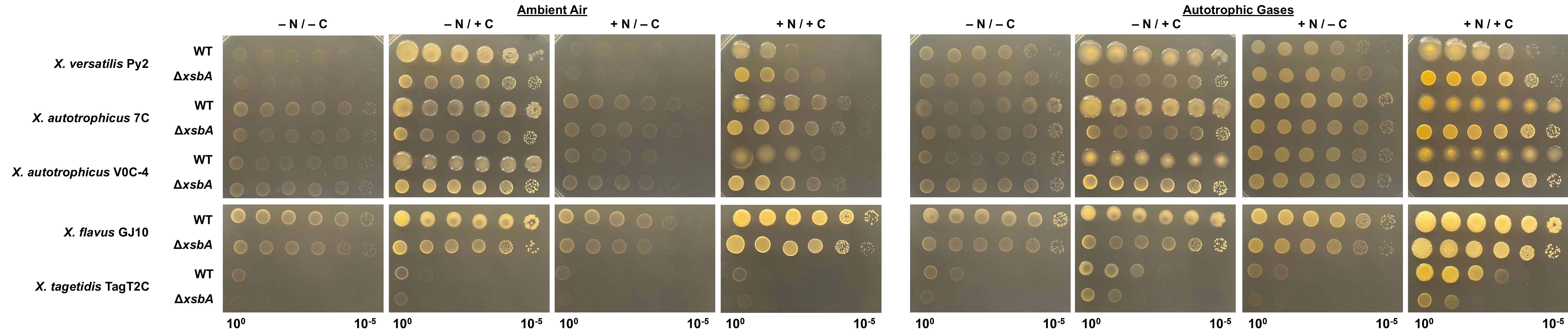

WMM + Bicarbonate

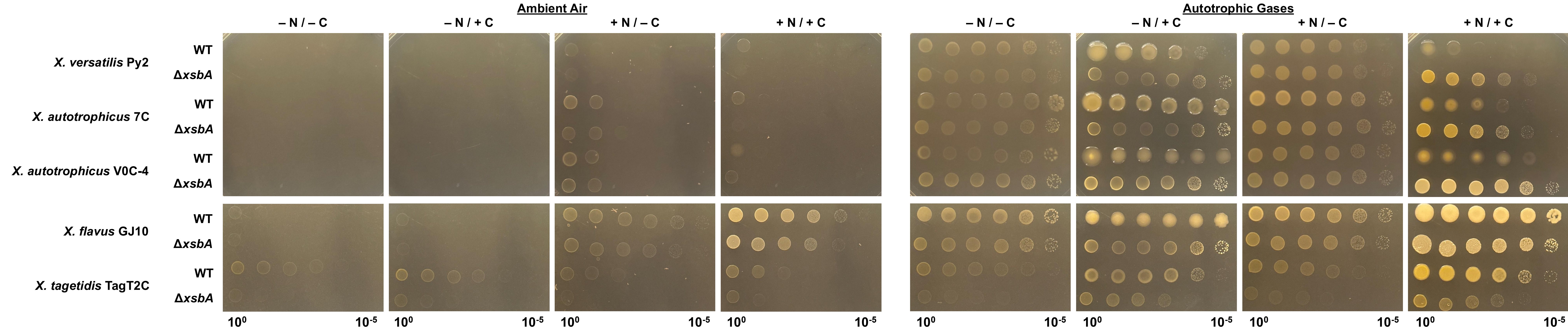

**Supplementary Table 9.** Xanthobacter strains used in this study. Abbreviations: ATCC = American Type Culture Collection; CCUG – Culture Collection University of Gotenburg; DSM(Z) – Deutsche Sammlung von Mikroorganismen und Zellkulturen GmbH/German Collection of Microorganisms and Cell Cultures GmbH; NCIMB – National Collection of Industrial, Food and Marine Bacteria. The asterisk indicates that generating the strain was attempted but not viable.

| Strain | Source | Used Experimental |  |  | NCBI Accessions Used for Bioinformatics |
| --- | --- | --- | --- | --- | --- |
|  |  | WT | + *xsba*<sup>7C</sup> | Δ*xsba* |  |
| *X. versatilis* Py2 | ATCC BAA-1158 | X | X | X | GCF_060823255.1, WP_506791320.1 |
| *X. versatilis* V1C-6B | Our Lab<br>(Available DSM 117963) |  |  |  | GCF_043193995.2, WP_394002759.1 |
| *X. oligotrophicus* 23A | DSM 685 | X |  |  | GCF_043194165.2, WP_188079529.1 |
| *X. autotrophicus* 7C | DSM 432 | X | X | X | GCF_043194295.1 (Table S1), GCF_043194295.2 (Rest), WP_138398082.1 |
| *X. autotrophicus* 7CSF | DSM 2267 | X | X |  | GCF_043194195.1, WP_394025923.1 |
| *X. autotrophicus* V0C-4 | Our Lab<br>(Available DSM 117702) | X | X | X | GCF_043194495.2, WP_394075003.1 |
| *X. autotrophicus* GZ29 | DSM 1393 | X |  |  | GCF_043194585.2, WP_394090543.1 |
| *X. autotrophicus* JW33 | DSM 1618 | X |  |  | MFG1284069.1 |
| *X. autotrophicus* 19/-lx | DSM 2009 | X |  |  | MFG1320789.1 |
| *X. autotrophicus* 124X | DSM 6696 | X |  |  | MFG1316047.1 |
| *X. autotrophicus* NCIMB11171 | NCIMB 11171 | X |  |  | MFG1279746.1 |
| *X. autotrophicus* CCUG44692 | CCUG 44692 | X |  |  | WP_138398082.1 |
| *Xanthobacter* sp. T101 | ATCC 700551 | X |  |  | GCF_043194595.2, WP_394111566.1 |
| *X. nonsaccharivorans* 14g | DSM 431 | X |  |  | GCF_043194755.2, WP_393978017.1 |
| *X. lutulentifluminis* V3C-3 | Our Lab<br>(Available DSM 117706) | X |  |  | GCF_043194675.2, WP_394114979.1 |
| *X. aminoxidans* 14a | DSM 15009 | X |  |  | GCF_043193655.2,<br>WP_250165692.1 |
| *X. flavus* 301 | DSM 338 | X |  |  | GCF_043193365.2,<br>WP_169124175.1 |
| *X. flavus* GJ10 | DSM 3874 | X | X | * | GCF_060823235.1,<br>WP_229643897.1 |
| *X. wiegelii* RH 10 | DSM 597 | X |  |  | GCF_043193855.2,<br>WP_394103047.1 |
| *X. cornucopiae* V4C-4 | Our Lab<br>(Available DSM 117704) | X |  |  | GCF_043195065.2 |
| *X. agilis* SA35 | DSM 3770 | X |  |  | GCF_043194985.2 |
| *X. sediminis* V8C-5 | Our Lab<br>(Available DSM 117708) | X |  |  | GCF_043195015.1 |
| *X. albus* V0C-6 | Our Lab<br>(Available DSM 117699) | X |  |  | GCF_043194895.2 |
| *X. variabilis* V4C-8 | Our Lab<br>(Available DSM 117713) | X |  |  | GCF_043831965.2 |
| *X. pseudotagetidis* KA | DSM 11602 | X |  |  | GCF_043194795.2,<br>WP_394016851.1 |
| *X. tagetidis* TagT2C | DSM 11105 | X | X | X | GCF_043194775.2,<br>WP_170168514.1 |
| *Xanthomonas campestris*<br>NRRL B-1459 | Genome Only (NCBI) |  |  |  | GCA_001372255.1,<br>WP_057672146.1 |
| *Ensifer/Rhizobium/Sinorhizobium*<br>*meliloti* 1021 | Genome Only (NCBI) |  |  |  | GCA_000006965.1,<br>WP_010975907.1 |
| *Escherichia coli* K-12 (MG1655) | Genome Only (NCBI) |  |  |  | NP_416566.1,<br>NP_310871.1,<br>NP_416531.4,<br>NP_310869.2,<br>NP_416564.4 |

**Supplementary Figure 10.** Serial dilutions of slime-producing and non-slime-producing *Xanthobacter* strains on 2xM1 or 2xM1 10% sucrose plates containing 0.08% w/v Congo Red. Plates were grown for 7 days at 30°C.

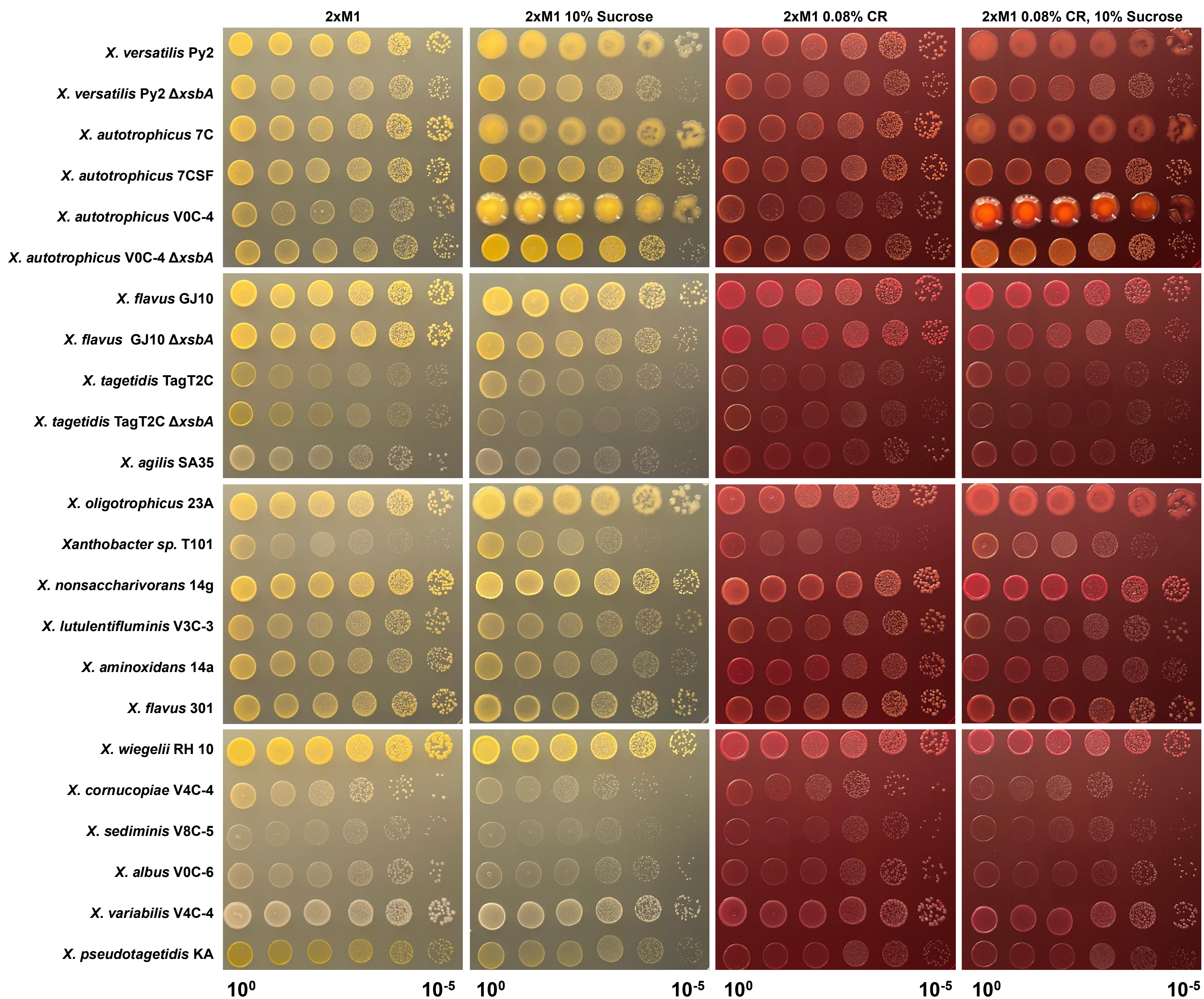
